# Layered helical order reveals assembly states in podosome actin networks

**DOI:** 10.64898/2026.07.24.740591

**Authors:** Jonathan Schneider, Javier Rey-Barroso, Thomas Drexler, Joel Valdivia Ortega, Stéphanie Balor, Dennis Ostlaender, Juergen M Plitzko, Wolfgang Baumeister, Renaud Poincloux, Marion Jasnin

## Abstract

How stochastic molecular events are converted into ordered cellular machines that perform mechanical work remains a central question in biology. Cryo-electron tomography enables visualization of macromolecular assemblies inside cells at molecular resolution, but static tomograms are generally interpreted as structural snapshots, with limited access to assembly state and temporal progression. Here, we show that cellular tomograms retain interpretable information about the assembly of a force-generating actin network. Using human macrophage podosomes as a physiologically relevant system, we combine spatial mapping of actin filaments (F-actin) and Arp2/3-mediated branch junctions, orientation analysis of deep-learning-based filament segmentations and Markov-chain modeling to infer actin-network assembly *in situ*. We identify favored membrane-directed actin polymerization and Arp2/3-mediated branching, establishing a local growth axis for podosome network organization. At the network scale, podosomes display layered helical order: filament-orientation classes recur at intervals of approximately 33 nm along the membrane-normal axis, forming a three-class cycle that returns to an approximately equivalent non-polar orientation. Markov modeling identifies preferred mother-to-daughter filament-orientation transitions and yields a stationary composition of branch-orientation states that closely matches that observed in structurally more advanced network regions. Together, these results link local actin nucleation, network-scale architecture and structural assembly state in a native cellular machine. Our study advances cryo-electron tomography from structural description toward temporal inference and 4D *in situ* structural biology.

## Introduction

Cryo-electron tomography (cryo-ET) has transformed structural cell biology by enabling protein complexes, membranes, organelles and cytoskeletal networks to be visualized at molecular resolution directly within near-native cellular environments (1). This creates an opportunity to connect molecular architecture with cellular function, particularly in large cytoskeletal assemblies whose activity depends not only on the structures of individual macro-molecules but also on their higher-order organization, spatial coordination and mechanical coupling (2). However, because cryo-ET captures dynamic cellular processes as static structural snapshots, extracting temporal information from such data remains a major challenge (3).

Recent studies have recovered dynamic or structural-state information from cellular tomograms by classifying conformational, compositional or maturation states of abundant molecular machines, including translating ribosomes and ribosomal subunits (4; 5; 6; 7), or by comparing nuclear-pore conformations across defined cellular conditions (8). F-actin networks pose a distinct challenge because they are compositionally and architecturally heterogeneous assemblies shaped by nucleators, boundary conditions and mechanical load (9). Coordinated actin polymerization drives membrane remodeling and force generation (9; 10; 11; 12), while forces can in turn remodel F-actin, altering filament conformation and mechanosensitive protein recognition (13). Understanding actin-network function therefore requires defining not only its molecular components but also the three-dimensional organization that emerges through assembly and mechanical remodeling.

To date, *in situ* analyses have resolved complementary aspects of actin-network organization. Filament polarity reveals the direction of F-actin growth at the level of individual filaments (14; 15; 16), whereas the orientation of Arp2/3-mediated branches, in which daughter filaments are nucleated from pre-existing mother filaments (17; 18), provides complementary information about daughter-filament growth relative to membrane-associated network geometry (19). *In situ* structures of Arp2/3-mediated branch junctions have also been obtained in actin waves and lamellipodia (19; 20). In highly ordered systems such as sarcomeres, cryo-electron tomography has resolved both whole-network filament organization and associated molecular components (15; 21; 22; 23). However, how local actin-assembly events collectively generate the three-dimensional architecture of a complete cellular network remains unresolved.

This gap is particularly important for Arp2/3-dependent branched actin networks, in which repeated nucleation builds higher-order organization. Such networks have been understood primarily through the dendritic nucleation model developed from studies of lamellipodia, where Arp2/3-mediated branching generates protrusive force at the leading edge of migrating cells (18). Yet many cellular actin structures differ from lamellipodia in their geometry, boundary conditions and mechanical output (24; 25). Whether Arp2/3-mediated nucleation follows common or structure-specific assembly principles in dense three-dimensional networks is therefore not known.

Macrophage podosomes are particularly well suited to address this question. They are submicrometric, actin-rich adhesion structures that exert nanonewton-scale forces on the extracellular environment and contribute to extracellular-matrix degradation, three-dimensional migration and phagocytosis (26; 27). These activities are central to macrophage behavior in tissues, where cells probe, remodel and migrate through complex extracellular environments during immune surveillance, inflammation and host defense (28). Podosome cores contain dense branched actin networks that depend on WASP-mediated activation of the Arp2/3 complex (29; 30). The cores are coupled to lateral filaments that connect them to an integrin-based adhesion ring and balance core protrusion with ring-associated traction (31). Podosome actin networks can also self-compress and behave as spring-loaded elastic materials that support force generation (32). Together, these properties make podosomes a physiologically relevant system for investigating how Arp2/3-mediated nucleation builds a native force-generating cytoskeletal machine.

Here, we tested whether spatial information encoded in static cryo-ET data can reveal how a native cellular actin network is assembled. We integrated F-actin polarity, Arp2/3-mediated branching and whole-network filament organization to infer how three-dimensional network architecture emerges and determine whether this architecture encodes assembly state.

## Results

### F-actin polarity reveals favored membrane-directed growth in podosomes

To establish the direction of network growth, we first mapped F-actin polarity to infer filament growth direction. Using cryo-ET data optimized for whole-podosome analysis (13 tomograms), we applied a subtomogram-averaging workflow that assigns the positions and orientations of individual actin monomers within filaments (15) (Methods and Supplementary Fig. S1). This generated a three-dimensional polarity map of the podosome actin network with monomer-level precision (Fig. 1).

**Figure 1.**
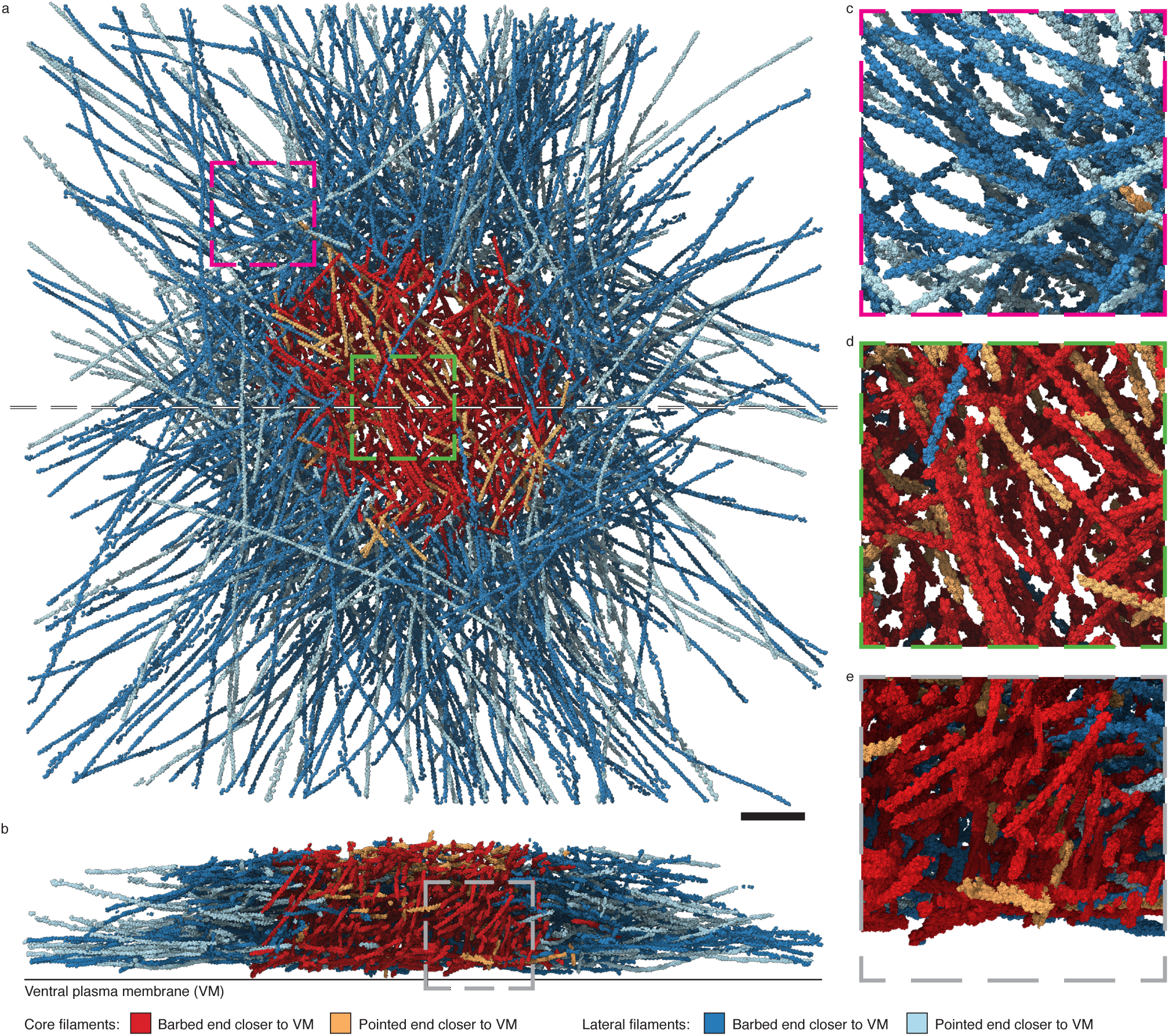
F-actin polarity reveals favored membrane-directed filament growth in podosomes. a,b, Top and side views of the polarity-resolved F-actin network in podosome #3. Colored boxes indicate regions enlarged in **c–e**. The dashed line in **a** indicates the position of the side-view section shown in **b**. Scale bar, 100 nm. **c–e,** Representative lateral (**c**) and core (**d,e**) filament regions. Core filaments are colored according to whether their barbed or pointed ends are closer to the ventral plasma membrane (red and orange, respectively), whereas lateral filaments are shown in dark and light blue, respectively.

We resolved the podosomal F-actin structure to 8.8 Å resolution and assigned polarity to 79 *±* 15% of filaments across 13 podosomes (Fig. 1 and Supplementary Figs. S1 and S2a). Actin filaments were classified as belonging to the core, characterized by upright filaments (32), or to the adhesion ring. In both regions, most filaments had their barbed ends directed toward the ventral plasma membrane (65 *±* 14% in the core and 66 *±* 6% in the adhesion ring; Fig. 1 and Supplementary Fig. S2a). Thus, F-actin growth was preferentially, although not exclusively, directed toward the ventral plasma membrane throughout the podosome.

We next examined how lateral filament polarity was organized relative to the podosome core. Pointed ends were concentrated around the inner core periphery and the core–adhesion ring interface (Supplementary Fig. S2b). Consistent with this spatial distribution, 70 ± 9% of lateral filaments were oriented outward, with their barbed ends positioned farther from the core than their pointed ends. Together, these polarity patterns indicate that lateral actin filaments form an outward-growing, membrane-directed network that extends from the podosome core across the adhesion ring.

### Arp2/3-dependent branching supports podosome architecture and force generation

We next examined how partial Arp2/3 inhibition affects podosome architecture and force generation. The Arp2/3 complex is enriched in podosome cores (Supplementary Fig. S3a,a’) and is required for podosome formation (29). Because complete inhibition prevents podosome formation, we partially inhibited Arp2/3 activity with 25 µM CK666 (Supplementary Fig. S3b–d). CK666 treatment reduced podosome number by 59 ± 18% and slowed the incorporation of actin–RFP into podosome cores by a factor of 1.7 (Supplementary Fig. S3e–g), consistent with reduced Arp2/3-dependent actin assembly.

Cryo-ET of CK666-treated cells revealed clusters of smaller podosomes, with a median core radius corresponding to 56% of the control value (207 nm versus 369 nm; Fig. 2a,b and Supplementary Fig. S3h,i). Partial Arp2/3 inhibition also significantly decreased the numbers of core and lateral filaments and their cumulative lengths (Supplementary Fig. S3j–m). Thus, Arp2/3-mediated nucleation contributes not only to core assembly but also to the actin network extending into the adhesion ring.

**Figure 2.**
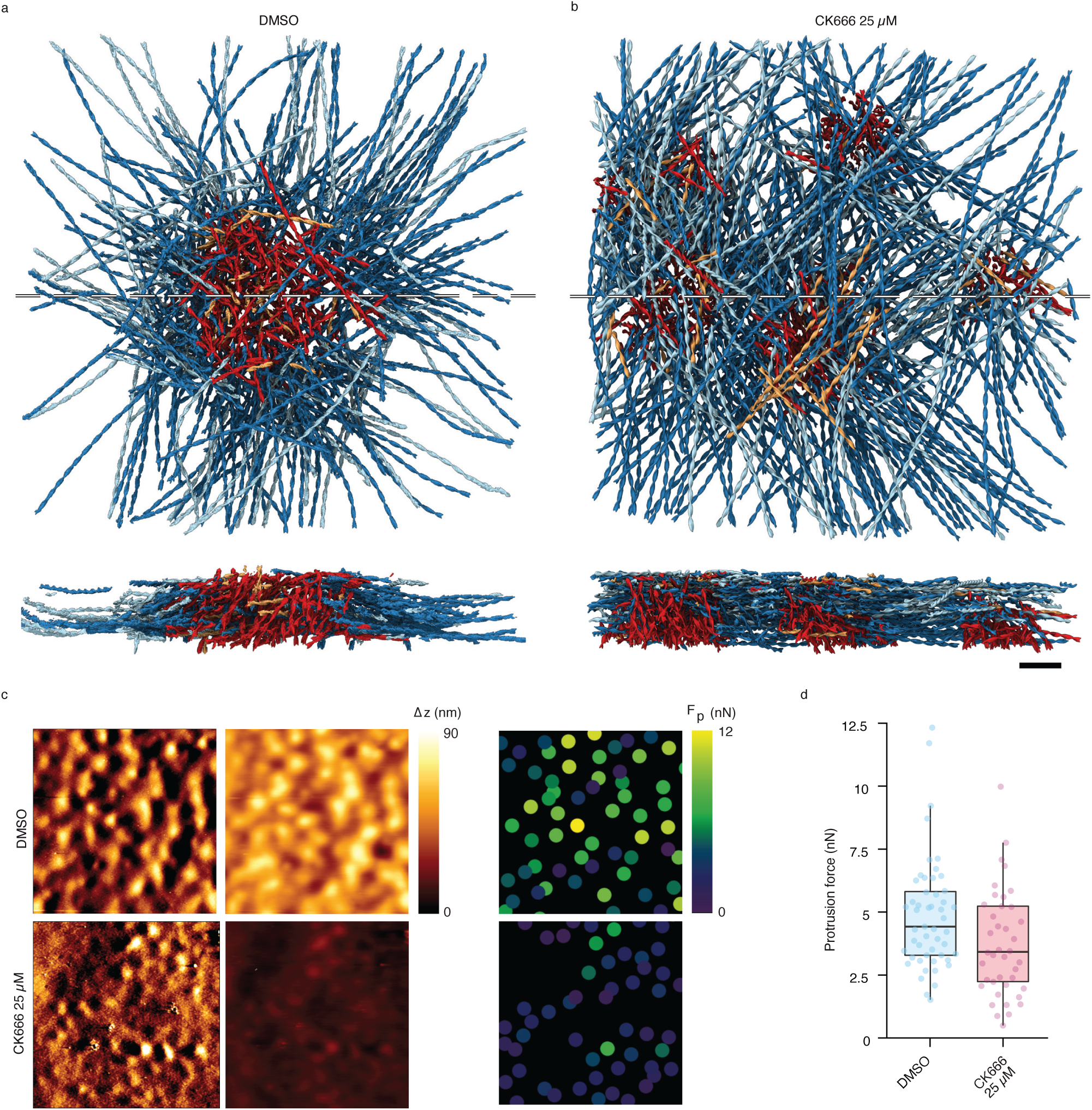
Arp2/3-dependent branching supports podosome architecture and force generation. a,b, Top and side views of polarity-resolved F-actin in a DMSO-treated podosome #5 (**a**) and a 25 *µ*M CK666-treated podosome cluster #2 (**b**). Dashed lines indicate the positions of the side-view sections. Filament colors follow the convention in Fig. 1. Scale bar, 100 nm. **c,** Representative vertical-deflection, deformation-height and protrusion-force maps under control and CK666-treated conditions. Scale bar, 10 *µ*m. **d,** Podosome protrusion forces. Each point represents the median force per cell; boxes show the median and interquartile range, with 10^th^–90^th^ percentile whiskers. *n* = 3 donors, 70 control cells and 57 CK666-treated cells, comprising more than 2,000 podosomes. Two-tailed Mann–Whitney test; \*\*\**P <* 0.0001.

To determine whether the structural changes induced by partial Arp2/3 inhibition were associated with altered mechanics, we measured protrusive forces using atomic force microscopy (33). CK666 treatment reduced the forces exerted on the substrate by 31 *±* 21% compared with control cells (Fig. 2c,d). Together, these results show that Arp2/3-dependent branching supports podosome architecture and protrusive force generation.

### Arp2/3-mediated branching couples podosome core growth to lateral network expansion

To determine how Arp2/3-mediated nucleation contributes to podosome core assembly and lateral filament growth, we analyzed cryo-ET data optimized for branch-junction detection (19 tomograms; see Methods). Branch junctions were identified by template matching (34), followed by subtomogram averaging (Supplementary Fig. S4a), and mapped within podosome cores. Because of sample-thickness constraints at higher magnification, this dataset primarily captured smaller podosomes and podosome clusters.

To validate the particle assignments and establish the molecular identity of the detected junctions, we classified and refined the branch-junction particles, yielding a 9.4 Å reconstruction of the Arp2/3-mediated branch junction (Fig. 3a and Supplementary Fig. S4b–d). The density resolved all Arp2/3 subunits together with adjacent actin monomers in the mother and daughter filaments, with the Arp2/3 complex density reaching approximately 8 Å resolution. Rigid-body fitting of an atomic model (PDB 8P94) showed close agreement with the density (Supplementary Fig. S4e).

**Figure 3.**
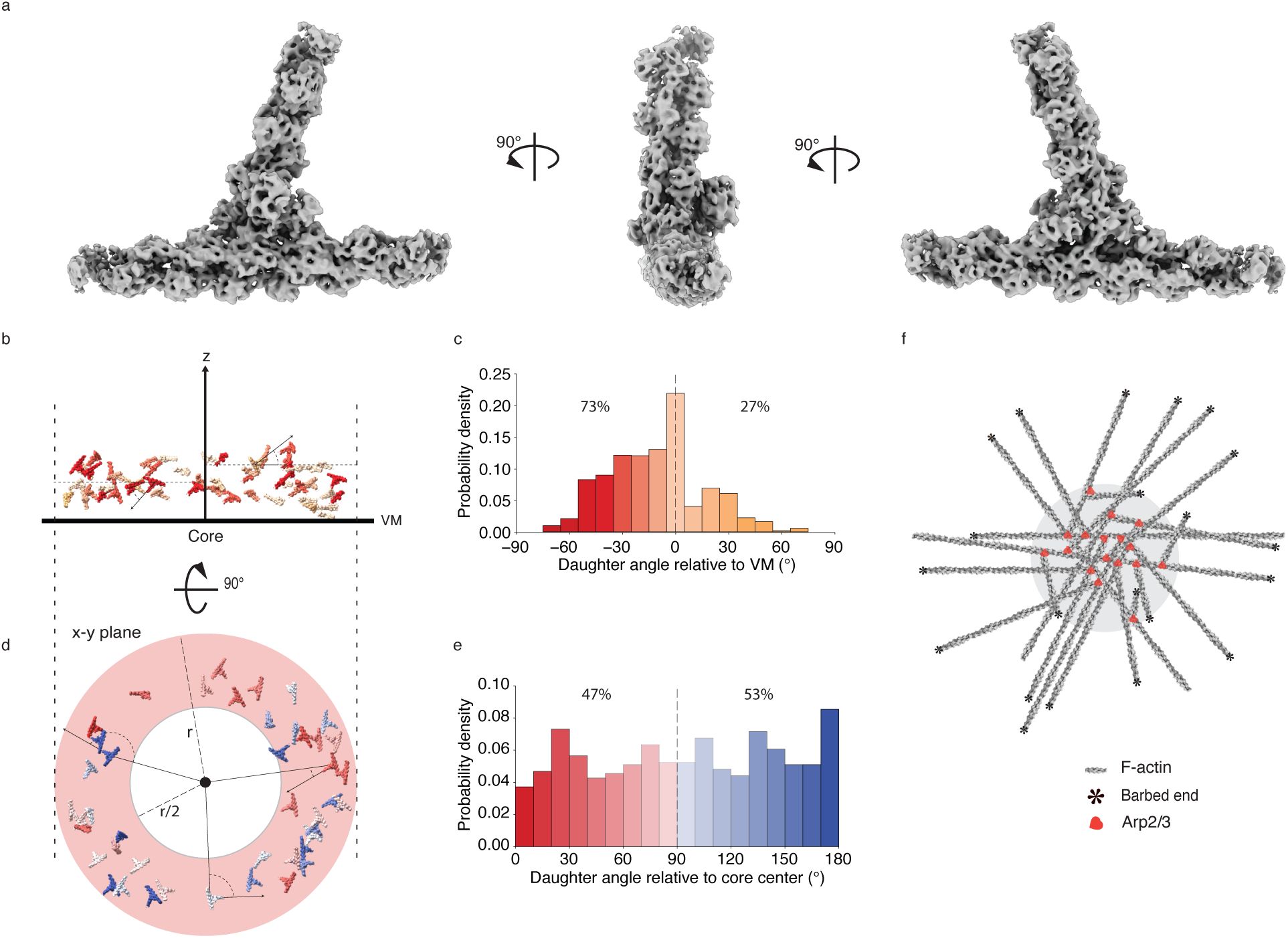
Arp2/3 branch-junction organization reveals directional filament growth in podosomes. a, Subtomogram average of the Arp2/3-mediated branch junction at 9.4 Å resolution, shown in three orientations. **b,c,** Side view of a representative podosome core #18 (**b**) and distribution of daughter-filament angles relative to the ventral plasma membrane (VM) across podosome cores (**c**). Colors indicate daughter-filament orientations toward or away from the VM. The analysis comprises 1,004 branch junctions from 11 cores in 7 tomograms. **d,e,** Top view of the same core (**d**) and distribution of daughter-filament angles relative to the core center within the shaded peripheral region (**e**), comprising 726 branch junctions from the same 11 cores. **f,** Schematic of divergent filament elongation. F-actin is shown in gray, with barbed ends marked by asterisks and Arp2/3 complexes in red.

We then used the refined branch positions and orientations to analyze the spatial organization of Arp2/3-mediated branching within podosome cores. Daughter filaments showed a strong bias toward the ventral plasma membrane, with 73% oriented toward the membrane, hereafter termed *downward* (Fig. 3b,c), consistent with the membrane-directed polarity of the podosome actin network (Figs. 1, 2a and Supplementary Fig. S2a). At the core periphery, daughter-filament orientations in the x–y plane were nearly evenly distributed relative to the podosome center, with 53% pointing outward and 47% inward (Fig. 3d,e). This branch-nucleation geometry is consistent with a mechanism by which Arp2/3-mediated nucleation can both expand the core and generate lateral filaments extending toward the podosome adhesion ring (Fig. 3f).

### Podosomes display layered-helical actin organization

We next asked whether the dense actin networks of podosomes exhibit higher-order organization beyond individual filaments and branch junctions. To address this question, we segmented F-actin in the whole-podosome and branch-junction datasets using ActinSeg, a deep-learning model for actin-network segmentation in cryo-ET data (35) (Fig. 4a,b and Supplementary Fig. S5a). These segmentations enabled us to map local filament orientations throughout each podosome network. After excluding upright core filaments, we identified three preferred filament-orientation classes, each represented at approximately equal abundance (Fig. 4c–f and Supplementary Fig. S5b,c).

**Figure 4.**
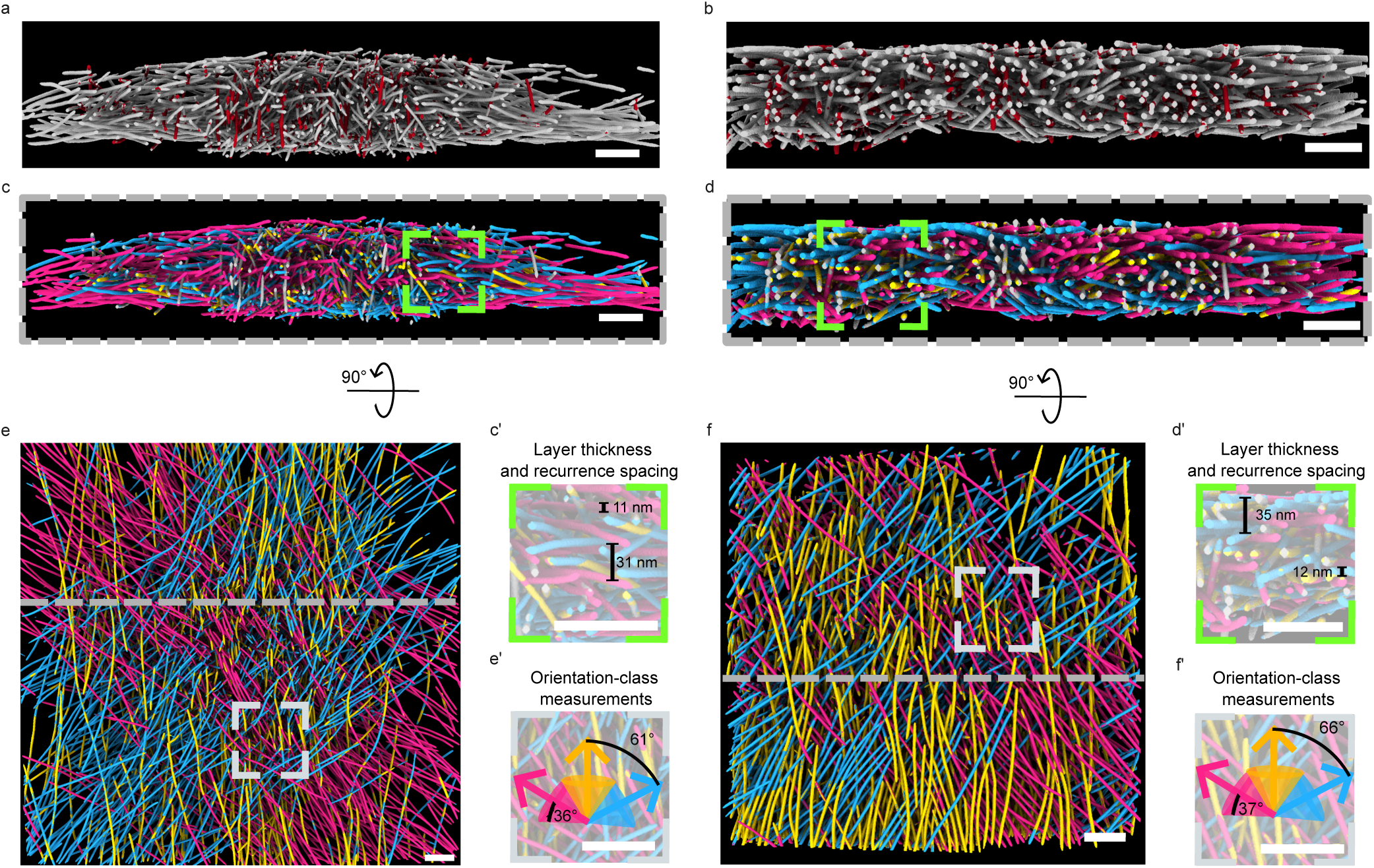
F-actin orientation analysis reveals layered helical order in podosomes. a,b, Representative side views of ActinSeg-derived F-actin segmentations from the whole-podosome (**a**) and branch-junction (**b**) datasets. Upright core filaments excluded from the orientation analysis are shown in red. **c,d,** Corresponding side-view orientation maps showing the three filament-orientation classes. **c’,d’,** Enlarged regions illustrating local layers of similarly oriented filaments and their axial recurrence. **e,f,** Corresponding top views. **e’,f’,** Enlarged regions showing the angular separation and within-class dispersion of the three orientation classes. Panels **a, c, c’, e, e’** show podosome #1; panels **b, d, d’, f, f’** show podosome #28. The dashed lines in **e** and **f** indicate the positions of the side-view sections shown in **a, c** and **b, d**, respectively. Scale bars, 100 nm.

Quantification showed that the three filament-orientation classes were separated by a characteristic angular interval. The mean angular separation between classes was 61 *±* 8°in the whole-podosome dataset and 66 *±* 13° in the branch-junction dataset. Thus, three-class filament-orientation organization was conserved across datasets acquired at different magnifications and sampling different podosome sizes. Within each class, filament orientations were distributed around a principal axis, with mean within-class angular dispersions of 36 *±* 23°and 37 *±* 23°in the whole-podosome and branch-junction datasets, respectively (Fig. 4e’,f’).

The three orientation classes were also spatially organized within the podosome network. Filaments assigned to the same class formed local same-orientation layers, defined as domains over which filament orientations remained approximately aligned. These layers did not form flat sheets spanning the entire podosome; instead, they formed local, undulating domains stacked along the membrane-normal axis within the continuous actin network. Along this axis, the three orientation classes alternated in a repeating sequence (Fig. 4c’,d’). Same-orientation layers had comparable mean thicknesses in the whole-podosome and branch-junction datasets, at 11.3 *±* 5.0 nm and 11.7 *±* 5.1 nm, respectively. Along the membrane-normal axis, each orientation class recurred at mean spacings of 30.9 *±* 6.6 nm and 34.9 *±* 5.2 nm, respectively.

We next quantified the angular closure of the three-class cycle. Because the orientations were treated as non-polar axes, a cumulative rotation of 180° returns the system to an equivalent orientation. In the whole-podosome dataset, the three-class cycle accumulated approximately 184°of rotation and therefore closed to within approximately 4°. Accordingly, the effective axial repeat of the layered organization was defined by the approximately 31 nm recurrence spacing of the same orientation class, a value numerically close to the long-pitch repeat of F-actin. In the branch-junction dataset, the accumulated rotation over one cycle was approximately 199°, corresponding to a residual offset of approximately 19°from the equivalent 180°orientation. The cycle therefore showed less complete angular closure in the branch-junction dataset. Whether this difference reflects the smaller podosomes preferentially sampled at higher magnification or an earlier structural assembly state remains to be determined.

Together, these results show that non-upright filaments in podosomes form a locally layered-helical organization along the membrane-normal axis. The three filament-orientation classes recur at intervals of approximately 33 nm, revealing coordination of filament orientations over length scales of tens of nanometers within the three-dimensional network.

### Markov modeling links branch-orientation transitions to reproducible podosome network organization

The layered-helical organization raised the possibility that reproducible network architecture could emerge from repeated local orientation transitions. Because Arp2/3-mediated branching generates daughter filaments from pre-existing mother filaments (17; 18), we asked whether preferred mother-to-daughter orientation transitions could contribute to this organization. We therefore analyzed these transitions across the podosome network (Methods). The analysis was not restricted to podosome cores because daughter-filament orientations relative to the ventral plasma membrane were similar across network regions, and inclusion of all detected branch junctions increased statistical power (Supplementary Fig. S6a).

To assess whether branch-orientation behavior was recurrent along the membrane-normal axis, we divided each podosome volume into successive 15-nm bands above the ventral plasma membrane and quantified daughter-filament orientation distributions within each band (Supplementary Fig. S6b,c). The recurrence of similar orientation distributions across successive bands suggested that network assembly could be described as repeated transitions among a common set of branch-orientation states.

We therefore modeled Arp2/3-mediated branching as a Markov process in which each branch event links a mother-filament orientation state to a daughter-filament orientation state with a defined transition probability. A three-component Gaussian mixture model applied to daughter-filament orientations relative to the ventral plasma membrane identified three dominant states, which we termed *downward*, *parallel* and *upward* (Methods and Fig. 5a). These membrane-referenced branch-orientation states are distinct from the non-polar orientation classes defining the layered architecture. Accordingly, we did not use the Markov model to reconstruct the layered pattern directly, but instead asked whether repeated local branching events could give rise to a reproducible network-wide orientation-state composition.

**Figure 5.**
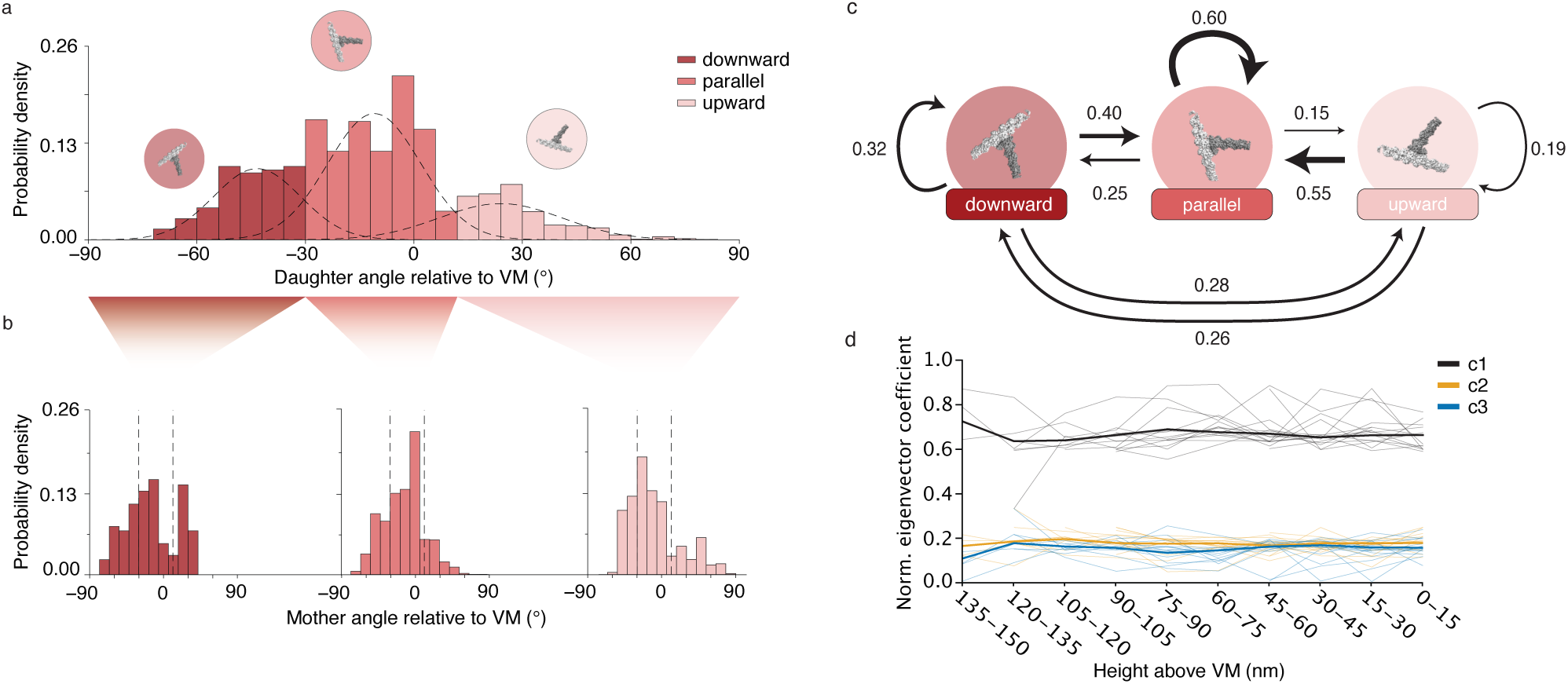
Markov modeling links branch-orientation transitions to reproducible podosome network organization. a, Distribution of daughter-filament angles relative to the ventral plasma membrane (VM) for 2,666 branch junctions from 17 tomograms. A three-component Gaussian mixture model defines *downward*, *parallel* and *upward* orientation states. **b,** Mother-filament angle distributions grouped by daughter-filament state. Dashed lines indicate state boundaries. **c,** Markov model linking the three branch-orientation states. Arrow thickness and labels indicate mother-to-daughter transition probabilities. **d,** Contributions of the transition-matrix eigenvectors to daughter-state distributions in successive 15-nm height bands above the VM, ordered from membrane-distal to membrane-proximal regions. *c*_1_ corresponds to the stationary eigenvector. Thick lines show means across tomograms; thin lines show individual tomograms.

For each daughter-filament state, we extracted the corresponding distribution of mother-filament orientations (Fig. 5b). These paired mother–daughter assignments were then used to construct a 3 *×* 3 transition matrix, in which each entry represents the probability that a mother filament in a given orientation state produces a *downward*, *parallel* or *upward* daughter filament (Fig. 5c).

Eigendecomposition of this matrix identified a unique stationary distribution comprising 27% *downward*, 54% *parallel* and 19% *upward* daughter-filament states. Because actin nucleation occurs at the ventral plasma membrane, membrane-proximal layers are assembled later in the growth sequence and were therefore considered to represent a more advanced network state. To test whether height-resolved compositions approached the stationary distribution, we expressed the composition in each height band as a linear combination of the normalized eigenvectors. The resulting coefficients showed convergence toward the stationary distribution in membrane-proximal regions (Fig. 5d).

We next quantified daughter-filament orientation states in the lower half of each podosome (Supplementary Fig. S6d). The resulting distribution, comprising 28% *downward*, 51% *parallel* and 21% *upward* states, closely matched the stationary distribution. Although both analyses were derived from the same network-wide dataset, this agreement indicates that the measured local transition probabilities are consistent with the state composition observed in more advanced network regions.

Together, these findings show that repeated Arp2/3-mediated nucleation events, although individually probabilistic, can generate a reproducible network-wide composition of branch-orientation states through defined transition probabilities.

### Podosome architecture provides an assembly pseudo-age

We next asked whether the progressive accumulation of recurrent orientation layers could provide a structural index of podosome assembly state. To quantify layer recurrence, we divided each tomogram laterally into a 3 *×* 3 grid and restricted the analysis to the central region. This reduced boundary effects from neighboring podosomes and enabled robust comparison of layered organization across tomograms. For each of the three filament-orientation classes, we calculated the spatial autocorrelation of its probability as a function of displacement along the membrane-normal axis (Fig. 6).

**Figure 6.**
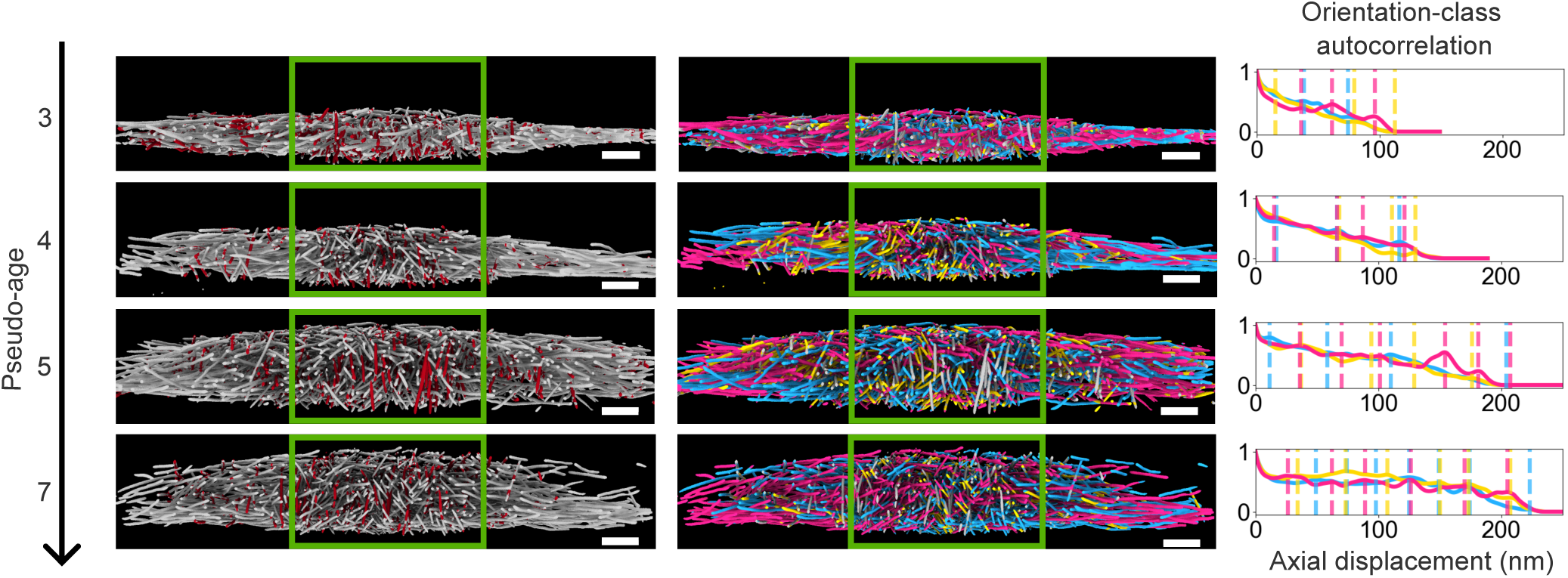
Pseudo-age estimation of podosomes based on layered structural order. Representative F-actin segmentations (left), filament-orientation maps (middle) and orientation-class autocorrelation profiles (right) for podosomes with increasing pseudo-age. Green boxes indicate the central regions used for analysis. Colored curves represent the three orientation classes, and vertical lines mark nonzero-offset autocorrelation maxima used for pseudo-age calculation. Podosomes correspond to #10, #4, #7 and #3, from top to bottom. Scale bars, 100 nm.

The resulting autocorrelation profiles decayed non-monotonically from 1 toward 0, with local maxima marking recurrent occurrences of the same filament-orientation class along the membrane-normal axis. The spacing between successive maxima therefore provided a measure of same-orientation layer recurrence. We defined the assembly pseudo-age of each podosome as 1 plus the rounded mean number of nonzero-offset local maxima detected across the three autocorrelation profiles. Podosomes with higher pseudo-ages generally extended farther along the membrane-normal axis, consistent with the inferred assembly progression. Because pseudo-age is based on orientation-layer recurrence rather than absolute podosome height, it is less directly affected by local membrane curvature or force-induced deformation. It should therefore be interpreted as a structural indicator of assembly progression rather than as a direct measure of chronological age.

## Discussion

Here, we show that human macrophage podosomes are organized as locally layered-helical actin networks whose architecture retains signatures of assembly state. By integrating F-actin polarity, Arp2/3-mediated branch geometry and whole-network filament orientation, we extend molecular-resolution actin analysis from individual structural features to an entire force-generating branched network. This framework links local nucleation events to reproducible three-dimensional architecture and enables structural-state inference from static cellular tomograms.

These results also extend the dendritic nucleation model established from quasi-two-dimensional lamellipodial networks, in which Arp2/3-mediated branching generates protrusive force against the leading edge (18; 36). Podosomes present a distinct geometric and mechanical problem: they are confined ventral structures in which core protrusion is coupled to radial adhesion and traction. At the core periphery, approximately half of the daughter filaments are oriented away from the core, consistent with their elongation across the adhesion ring and with a divergence–elongation mechanism for lateral network expansion (Fig. 3f). This geometry resembles principles observed in reconstituted and simulated branched actin networks, in which spatially confined Arp2/3-activating regions of comparable dimensions can guide lateral actin organization through nucleation geometry, filament mechanics and local filament interactions (37).

The divergence–elongation mechanism identified here places podosomes within a broader spectrum of Arp2/3-dependent actin architectures. It contrasts with the convergence–elongation model proposed for filopodia, in which filaments arise through convergence and bundling of a branched lamellipodial network (38). Podosomes also differ from ventral traveling actin waves, in which membrane-directed Arp2/3-mediated branching forms clustered, tent-like structures within a propagating network (19). Across these systems, daughter-filament growth is preferentially oriented toward membrane-associated regions where nucleation-promoting factors activate the Arp2/3 complex. This shared orientation bias suggests that interactions among Arp2/3, mother filaments and membrane-associated nucleation-promoting factors may favor particular branch orientations. However, the resulting architecture depends on whether the Arp2/3-activating region is planar and protrusive, propagating or geometrically confined. Arp2/3-mediated branching can therefore be tuned by nucleation-promoting context, spatial confinement, membrane geometry and mechanical output to generate distinct cellular actin architectures.

The layered helical architecture may also be influenced by molecular-scale properties intrinsic to F-actin. The same-orientation spacing in podosomes, approximately 33 nm, is close to the 35.9-nm long-pitch repeat of F-actin (39). A similar correspondence has been observed in lamellipodia, where closely spaced branch sites on the same filament were frequently separated by multiples of the actin helical repeat (40). Although the basis of this correspondence remains to be determined, it raises the possibility that the helical geometry of F-actin may influence the accessibility or probability of Arp2/3-mediated branching at particular positions on the filament surface, for example by modulating the exposure of branch sites to membrane-associated Arp2/3 activators. Podosome architecture may therefore emerge from the combined effects of membrane-associated Arp2/3 activation, geometric confinement, filament polarity and the intrinsic molecular geometry of F-actin.

In addition to these geometric and molecular factors, our results suggest that Arp2/3-dependent network organization is shaped by probabilistic transitions between branch-orientation states. The canonical 70°branch angle defines individual nucleation geometry but does not explain how repeated branching produces a reproducible network. Our transition analysis suggests that reproducible branch-state composition can emerge statistically from preferred mother-to-daughter transitions, with the stationary distribution providing a quantitative descriptor of the composition associated with structurally more advanced network regions.

Transition matrices may provide a common framework for comparing how Arp2/3-mediated branching is tuned across distinct cellular actin architectures. The *downward*, *parallel* and *upward* membrane-referenced orientation states used here were previously applied descriptively to ventral actin waves (19), but transitions between these states were not modeled in that study. Applying the Markov framework across systems could therefore test whether distinct network geometries arise through changes in the relative probabilities of the same elementary branch-orientation transitions. Although the Markov model does not capture longer-range history, filament interactions or mechanical feedback, it provides a compact quantitative description of how local branch-orientation biases relate to network-scale state composition.

This architectural framework also has a temporal implication. By quantifying the recurrence of same-orientation layers, we positioned individual podosomes along an inferred assembly trajectory. We refer to this structurally inferred state as assembly pseudo-age. Conceptually analogous to growth rings, recurrent orientation layers preserve structural information about assembly progression (41). This metric orders individual podosomes by structural assembly state without equating structural progression with chronological age.

More broadly, these findings expand how static cryo-ET data can be interpreted. Podosomes provide an initial case study because their networks are built through repeated Arp2/3-mediated nucleation, but related approaches may be applicable to endocytic actin networks and other cellular assemblies generated through recurring local transitions. Linking molecular transitions to higher-order architecture may therefore allow structural snapshots to reveal not only how cellular machines are organized, but also where they lie along their assembly trajectories.

## Methods

### Differentiation and culture of primary monocyte-derived macrophages

Human peripheral blood mononuclear cells were isolated from the blood of healthy donors by centrifugation through Ficoll-Paque Plus (Dutscher). Cells were resuspended in cold phosphate-buffered saline (PBS), pH 7.4, supplemented with 2 mM EDTA and 0.5% heat-inactivated fetal calf serum (FCS). CD14-positive monocytes were isolated by magnetic sorting using anti-CD14 magnetic microbeads (Miltenyi Biotec, 130-050-201). Monocytes were seeded on glass coverslips at 1.5 *×* 10^6^ cells per well in six-well plates containing RPMI 1640 medium (Invitrogen) without FCS. After 1 h at 37 *^◦^*C in a humidified atmosphere containing 5% CO_2_, the medium was replaced with RPMI 1640 supplemented with 10% FCS and 20 ng ml*^−^*^1^ macrophage colony-stimulating factor (M-CSF; PeproTech). Cells were differentiated for 7 days. For experiments, macrophages were detached using trypsin–EDTA (Fisher Scientific) and collected by centrifugation at 320 *×g* for 10 min.

For partial inhibition of Arp2/3 complex activity, macrophages were treated with CK666 at the indicated concentrations and incubation times. For cryo-ET experiments, cells were treated with 25 µM CK666 for 30 min before vitrification. Control cells were treated with 0.1% DMSO.

### Immunofluorescence

Macrophages were plated on glass coverslips for 3 h before fixation. Unless otherwise stated, cells were fixed for 10 min at room temperature in PBS containing 3.7% paraformaldehyde and 15 mM sucrose. When indicated, cells were unroofed before fixation by incubation for 30 s at room temperature in distilled water containing Halt protease and phosphatase inhibitor cocktail (Thermo Fisher Scientific, 78440) and 10 µg ml*^−^*^1^ phalloidin (Sigma-Aldrich, P2141), followed by 10 washes. After fixation, cells were permeabilized for 10 min in PBS containing 0.3% Triton X-100 and blocked in PBS containing 0.3% Triton X-100 and 1% bovine serum albumin (BSA). Samples were incubated with primary antibodies for 2 h at room temperature, followed by incubation for 1 h at room temperature with Alexa Fluor 546–phalloidin (Thermo Fisher Scientific, A22283; 1:300) and Alexa Fluor 488–conjugated secondary antibodies (Thermo Fisher Scientific, A32731; 1:500) to label F-actin and associated proteins, respectively.

### Fluorescence imaging and podosome count and size analysis

For podosome count and size measurements, unroofed macrophages were stained for F-actin with Alexa Fluor 546–phalloidin and for vinculin using Alexa Fluor 488–conjugated secondary antibodies (Molecular Probes, M4409; 1:500). Podosome number was quantified from F-actin-stained podosome cores using the *Find Maxima* function in ImageJ. More than 200 cells from at least three independent experiments were analyzed for each condition. Statistical comparisons were performed using the non-parametric Kruskal–Wallis test followed by Dunn’s multiple-comparison test. Statistical significance was defined as *P <* 0.001.

For podosome size measurements, samples were imaged in nine focal planes separated by 100 nm using a 100*×*/1.45 numerical-aperture objective on a Nikon Eclipse Ti-E microscope equipped with an Andor Neo sCMOS camera operated in high-dynamic-range and global-shutter modes. Image stacks were processed by nonlinear deconvolution using DeconvolutionLab in ImageJ, as described previously (31). The point-spread function used for deconvolution was obtained by averaging 30 experimentally measured point-spread functions. Podosome size was measured automatically from paired deconvolved F-actin and vinculin images using a dedicated ImageJ macro (33). Briefly, the center of each actin core was determined from local actin-intensity maxima weighted by the surrounding grey values. Radial vinculin-intensity profiles were used to measure the adhesion-ring radius, defined as the distance from the actin-core center to the peak vinculin intensity. For each podosome, the adhesion-ring radius was calculated as the median of eight radial measurements acquired at 45*^◦^* intervals. The donor-specific mean adhesion-ring radii used in the protrusion-force calculations are reported in Table 1. These fluorescence-derived adhesion-ring radii are distinct from the actin-core radii measured in the cryo-ET datasets.

**Table 1.** Donor-specific mean podosome adhesion-ring radii used for protrusion-force calculations. Percentage reduction was calculated relative to the donor-matched DMSO control. The summary row reports the mean *±* standard deviation (SD) across donors (*n* = 3).

| Donor ID | Mean adhesion-ring radius (nm) |  |  |
| --- | --- | --- | --- |
|  | DMSO control | CK666-treated | Reduction (%) |
| 37 | 486 | 403 | 17.1 |
| 235 | 497 | 435 | 12.5 |
| 818 | 404 | 389 | 3.7 |
| **Mean ± SD** | **462 ± 41** | **409 ± 19** | **11.0 ± 5.5** |

### Protrusion force microscopy

Protrusion force microscopy was performed as described previously (26; 33). Briefly, atomic force microscopy (AFM) measurements were performed using silicon nitride cantilevers (MLCT-AUHW, Veeco Instruments) with a nominal spring constant of 0.01 N m*^−^*^1^, mounted on a NanoWizard III AFM (JPK Instruments) coupled to an inverted optical microscope (Axiovert 200, Carl Zeiss). Cantilever sensitivity and spring constant were calibrated before each experiment using the thermal-noise method implemented in the JPK Instruments software.

Formvar-coated grids were prepared by dipping ethanol-cleaned glass slides into 0.5% (w/v) Formvar in ethylene dichloride (Electron Microscopy Sciences). After drying, the Formvar film was detached from the slides by contact with water and allowed to float on the water surface. Acetone-washed 200-mesh nickel grids (Electron Microscopy Sciences) were placed onto the floating film, collected on glass slides and air-dried. Formvar-film thickness was measured by AFM at the edge of the film remaining on the glass slide after removal of the grids.

Living macrophages were plated on Formvar-coated grids placed in a temperature-controlled chamber (Petri dish heater, JPK Instruments). The culture medium was supplemented with 10 mM HEPES, pH 7.4 (Sigma-Aldrich). Where indicated, cells were treated with 25 *µ*M CK666 for 30 min before force measurements. Images were acquired in liquid in contact mode using scanning forces below 1 nN, image sizes of 256 *×* 256 or 512 *×* 512 pixels and a line rate of approximately 2 Hz.

Forces exerted by individual podosomes were derived from the topographical profiles of podosome-induced protrusions as described previously (26; 33). Briefly, protrusion profiles were extracted from AFM images using an ImageJ macro. Deformation height was determined from the protrusion profile and adhesion-ring radius and converted into force using the established Formvar-deformation model. The biaxial Young’s modulus of Formvar, *E/*(1 *−ν*^2^), was 2.3 GPa (26), and *C*_0_ *≈* 2.7 was used as the geometric coefficient derived from numerical simulations (33). Film thickness and adhesion-ring radius were measured for each experimental series by AFM and immunofluorescence, respectively, using the donor-specific adhesion-ring radii reported in Table 1. Each cell was assigned the median force measured across its podosomes.

The analysis included 70 control cells and 57 CK666-treated cells from three independent experiments. Data distributions deviated significantly from normality according to the D’Agostino–Pearson test. The two groups were therefore compared using a two-tailed Mann–Whitney test. Differences were considered statistically significant at *P <* 0.0001.

### Fluorescence recovery after photobleaching

Fluorescence recovery after photobleaching (FRAP) was performed using a protocol established previously (26). Primary human macrophages were transfected with mRFP–actin (kindly provided by S. Linder) and plated in glass-bottom chambers (Lab-Tek) 24 h before imaging. FRAP experiments were performed on an LSM 710 confocal microscope (Zeiss) using the dedicated FRAP module in the ZEN software. The microscope was equipped with a 40*×*/1.3 numerical-aperture oil-immersion objective and an environmental chamber maintained at 37 *^◦^*C and 5% CO_2_.

The 561-nm laser line was used for both imaging and photobleaching. Images were acquired every 5 s with a pixel dwell time of 9 µs. Individual podosomes were defined using circular regions of interest with a diameter of 1.05 µm and were bleached sequentially to produce a 50 *±* 10% decrease in fluorescence intensity. Fluorescence-recovery curves and control curves obtained from non-bleached regions were measured in ImageJ, exported to Excel and fitted using the Soumpasis equation (42). Recovery half-times (*t*_1_*_/_*_2_) were calculated from the fitted curves.

The analysis included 36 control cells and 24 CK666-treated cells from three independent experiments. The two groups were compared using a two-tailed Mann–Whitney test. The difference was statistically significant (*P* = 0.0027).

### Cell unroofing and vitrification on EM grids

Gold EM grids bearing Quantifoil R 1/4 holey carbon or SiO_2_ support films (Quantifoil Micro Tools, Großlöbichau, Germany) were glow-discharged. Macrophages were resuspended together with fiducial markers and seeded onto the EM grids. Cells were incubated for 2 h at 37 *^◦^*C to allow adhesion, resulting in approximately three to four cells per grid square.

Cells were unroofed for 30 s using distilled water containing complete protease inhibitor cocktail (Roche) and 10 µg ml*^−^*^1^ phalloidin (Sigma-Aldrich, P2141). For vitrification, 3.4 µl of unroofing solution was applied to each grid. Grids were plunge-frozen in liquid ethane cooled by liquid nitrogen using an EM GP automatic plunge freezer (Leica) operated at 95% humidity. Grids were blotted from the reverse side for 10 or 11 s using Whatman No. 1 filter paper.

### Cryogenic tilt-series acquisition

Two complementary cryo-ET datasets were analyzed: a lower-magnification dataset capturing complete podosomes for polarity and whole-network analyses, and a higher-magnification dataset optimized for Arp2/3 branch-junction detection while also enabling filament-orientation analysis.

The whole-podosome dataset was acquired using SerialEM on a Titan Krios G2 transmission electron microscope (Thermo Fisher Scientific) operated at 300 kV and equipped with a field-emission gun, a post-column energy filter (Gatan) and a 4k *×* 4k K2 Summit direct electron detector (Gatan). Tilt series were recorded in counting mode at a nominal magnification of 42, 000*×*, corresponding to a pixel size of 3.42 Å. Tilt series were acquired over an angular range of *−*60*^◦^*to +60*^◦^* in 2*^◦^* increments using a dose-symmetric tilt scheme, with a target defocus range of *−*3 to *−*5 µm and a total electron dose of 150–200 e*^−^* Å*^−^*^2^.

The branch-junction dataset was acquired using Tomography 5 software on a Krios G4 transmission electron microscope (Thermo Fisher Scientific) operated at 300 kV and equipped with a cold field-emission gun, a Selectris X energy filter and a Falcon 4i direct electron detector. Tilt series were recorded in counting mode at a nominal magnification of 64, 000*×*, corresponding to a pixel size of 2.06 Å. Tilt series were acquired over an angular range of *−*54*^◦^*to +54*^◦^* in 2*^◦^* increments, with a target defocus range of *−*2.5 to *−*4.5 µm and a total electron dose of 190–200 e*^−^* Å*^−^*^2^.

### Tomogram reconstruction

For the whole-podosome dataset used for F-actin polarity and network-scale analyses, tilt-series preprocessing was performed using TOMOMAN (43) (Supplementary Fig. S1a). Movie frames were aligned using MotionCor2 (44) and dose-filtered according to the cumulative electron dose using exposure-dependent attenuation functions and critical exposure constants (45). Tomograms were reconstructed by weighted back-projection in IMOD (46), yielding a final reconstructed pixel size of 13.68 Å. To enhance contrast, tomograms were denoised using Cryo-CARE (47) and corrected for the missing wedge using IsoNet (48). In total, the whole-podosome analysis included 13 tomograms of untreated unroofed cells, each containing one podosome, and three tomograms of CK666-treated unroofed cells containing 12 podosomes in total.

For the branch-junction dataset used for Arp2/3 detection and branch-orientation analysis, tilt series were processed using AreTomo3 version 2.1.10 (49) (Supplementary Fig. S4a), yielding a final reconstructed pixel size of 8.24 Å. In total, 19 tomograms of untreated unroofed cells containing podosomes were included in the Arp2/3 branch-junction analysis. AreTomo3 was installed and configured by SBGrid (50).

### F-actin segmentation

For F-actin polarity analysis, actin filaments were segmented from binned tomograms using the filament-tracing workflow described previously (32). Briefly, tomograms were first subjected to non-local means filtering in Amira (Thermo Fisher Scientific). Actin filaments were then traced using an automated segmentation algorithm based on a generic filament template (51), with a filament diameter of 8 nm and a template length of 42 nm. Filamentous structures shorter than 60 nm were excluded to reduce background detections. Filament coordinates were exported from Amira and used for downstream analyses in MATLAB (The MathWorks).

For filament-orientation and layered-architecture analyses, F-actin was segmented using ActinSeg (35), a deep-learning model developed for actin-network segmentation in cryo-ET data. The model was trained on published datasets of podosomes (32), actin waves (19) and sarcomeres (15). ActinSeg was applied to the whole-podosome dataset, comprising 13 tomograms acquired at a nominal magnification of 42, 000*×*, and to the branch-junction dataset, comprising 19 tomograms acquired at a nominal magnification of 64, 000*×*. The resulting segmentations were used to extract local filament orientations throughout the podosome volumes and to analyze filament-orientation classes, same-orientation layers and higher-order podosome architecture.

### F-actin subtomogram analysis and polarity assignment

#### Filament resampling

Filament traces obtained from the Amira segmentations were resampled using a custom MATLAB script (https://github.com/jasnin-lab/podosomes.git). Coordinates along each filament were sampled at equally spaced intervals corresponding to half the axial spacing between actin monomers, i.e. 1.38 nm. At each sampling position, the local filament centerline was aligned with the *z*-axis, such that the angle *φ* described rotation about the filament axis. Because this rotation could not be determined directly from the tomograms, *φ* was sampled in 30*^◦^* increments. Each resampled point was assigned the unique identifier of its parent filament.

#### Generation of an F-actin reference and polarity assignment

F-actin polarity analysis was performed using the workflow described previously (15).

An initial F-actin reference was generated from 57 filaments longer than 550 nm. From these filaments, 27,550 subtomograms were extracted at 2*×* binning, corresponding to a pixel size of 6.84 Å, using a box size of 128 pixels. F-actin subtomogram averaging was performed in STOPGAP version 0.7.0 (52). Multiclass alignment was initiated from a cylindrical reference.

Sixteen filament averages displaying clear helical features were selected to generate the polarity reference. From these filaments, 8,193 unbinned subtomograms were extracted and refined separately. The best-resolved filament reconstruction and a copy rotated by 180*^◦^* were used as references for supervised polarity classification. Particles assigned to the opposite polarity were rotated by 180*^◦^* and combined with those assigned to the reference polarity. Joint averaging and local alignment were then performed to generate an improved, uniformly oriented F-actin reference. This reference was used in subsequent rounds of polarity assignment, including the reassignment of previously unclassified particles. A final refinement was performed after rotating all particles into a common polarity.

Filament polarity was assigned by multi-reference alignment in STOPGAP (52). Subtomograms were extracted at 2*×* binning, corresponding to a pixel size of 6.84 Å, using a box size of 64 pixels. Initial polarity assignment was performed on five tomograms using the reference generated above. Alignment comprised global in-plane angular searches, local out-of-plane cone searches and additional in-plane refinement, with exposure, CTF, cosine and score weighting enabled. An FFT threshold of 800 and a low-pass filter of 24 Å were applied.

For the remaining tomograms, the refined F-actin reconstruction was used as the reference. To reduce computational cost, a single alignment iteration was performed, with a maximum of 100 particles sampled per filament. A low-pass filter of 15 Å was applied. For each filament, alignment scores for the two possible polarities were grouped according to orientation and compared using a paired Student’s *t*-test. Filaments with *P <* 0.05 were assigned to the polarity with the higher mean alignment score. Their particles were then rotated to generate a consistently oriented particle list. A second round of polarity assignment was performed for filaments without significant score differences, and newly assigned filaments were added to the final dataset. Polarity-assignment success was quantified for each tomogram as the fraction of filaments assigned a polarity among all analyzed filaments.

#### F-actin subtomogram averaging

Five tomograms containing 1,643,230 particles were used for initial subtomogram averaging. Subtomograms were extracted at 2*×* binning using a box size of 64 pixels and aligned against a de novo reference low-pass filtered to 40 Å. Global in-plane alignment was followed by local cone search and subsequent local refinement of all Euler angles. Translational shifts along the filament axis were restricted to *±* 13.8 Å to prevent convergence onto non-neighboring actin-monomer positions. Distance-based cleaning was applied independently to each filament using a threshold of 20.7 Å, retaining one particle per actin monomer.

Due to memory limitations, only four tomograms containing 445,636 particles were selected for final refinement. Particles were divided into half-sets per filament. Local angular refinement was performed with progressively decreasing search ranges, followed by re-extraction at full resolution using a box size of 128 pixels. Ten additional alignment rounds were performed using full-cone searches over all Euler angles, with angular search ranges progressively reduced from *±* 8*^◦^* to *±* 1*^◦^*and a final angular step of 0.25*^◦^*. The refined average was used to generate a binary body mask in RELION 5 (53). This mask was multiplied by a softened spherical mask to reduce artifacts along the filament axis. The final resolution was determined by Fourier shell correlation using the resulting mask. A *B* factor of *−*1000 Å^2^ was applied to sharpen the final map, which reached a resolution of 8.8 Å.

### Podosome geometric analysis

#### Podosome core segmentation and diameter measurement

For each podosome, the core region was manually defined in a central tomographic slice. A boundary polygon was drawn in IMOD (46) to encompass the main core-filament population, including most upright filaments. Filaments were classified as core filaments when more than two-thirds of their traced length lay within this boundary. Filaments outside the core boundary were classified as non-core filaments and evaluated for inclusion in the lateral-filament analysis described below. The podosome center was defined as the arithmetic mean of all particle coordinates belonging to core filaments.

Core diameter was quantified by fitting an ellipse to the polygon vertices defining the core boundary and calculating the mean of the major-and minor-axis lengths. The actin-core radius reported in the Results was defined as one-half of this mean fitted diameter.

#### Ventral plasma membrane tracing and membrane-relative filament orientation

The ventral plasma membrane was manually segmented in IMOD (46) by placing splines along the membrane contour across successive tomographic slices. The spline coordinates were used to fit a polynomial surface to the membrane in MATLAB. The interpolated surface was then converted into a binary, single-voxel-thick representation of the ventral plasma membrane.

For each podosome, the local ventral plasma membrane normal was determined from the membrane region underlying the podosome core. Coordinates defining the core boundary were projected onto the fitted membrane surface to generate a set of three-dimensional membrane points. A plane was fitted to these points using MATLAB functions, and its normal vector was used as the reference direction for membrane-relative orientation analysis.

For each filament with assigned polarity, the sampling point closest to the ventral plasma membrane was identified. A unit vector oriented along the mean local filament direction was applied from this point, and the distance between the displaced point and the fitted membrane surface was compared with the original distance. Filaments were classified as barbed-end directed toward the membrane when the displaced point was closer to the membrane surface, and as pointed-end directed toward the membrane when it was farther away.

#### Lateral-filament analysis

Lateral filaments were defined as non-core filaments located within a radial distance of 1.25 times the core radius from the podosome center. For each lateral filament, the mean filament orientation was calculated from the Euler angles along the filament trace, and the filament center was defined as the arithmetic mean of its particle coordinates. A unit vector aligned with the mean filament direction was applied from the filament center, and the distance of the displaced point from the podosome center was compared with the original filament-center distance. Filaments were classified as radially outward when the displaced point was farther from the podosome center and as radially inward when it was closer.

For CK666-treated samples containing multiple podosome cores within the same tomogram, each core was analyzed independently. Consequently, filaments located within 1.25 core radii of more than one core could contribute to the analysis of multiple podosomes.

### Arp2/3-mediated branch-junction analysis

#### Branch-junction template matching and subtomogram averaging

Template matching was performed on the 19 tomograms of the branch-junction cryo-ET dataset acquired at 64, 000*×* magnification using `pytom_match_pick` from PyTom (34). The initial template was generated from a previously reported Arp2/3-mediated branch-junction structure (EMDB-15136), low-pass filtered to 16 Å and resampled to the tomogram pixel size of 8.24 Å per pixel using `create_template.py`. A matching mask was generated using a custom Python script and consisted of two cylindrical volumes oriented at an angle of 70*^◦^* relative to each other. This mask enclosed the branch-junction geometry while excluding filament ends. The template and mask were matched in pixel size and box dimensions.

Template matching was performed using the parameters listed in Supplementary Fig. S4. Receiver operating characteristic curves generated with `pytom_estimate_roc_curves.py` were used to determine a particle-extraction threshold for each tomogram. Particle coordinates were extracted using the corresponding tomogram-specific threshold. Metadata generation and conversion to RELION 5 format (53) were performed using `aretomo3torelion5.py` (49). Particles were subsequently processed in RELION 5 according to the workflow shown in Supplementary Fig. S4.

Template matching and subtomogram averaging were performed iteratively. In the first iteration, the external Arp2/3 branch-junction template was matched against five tomograms, yielding 648 particles. Particles were extracted at bin 4, corresponding to a pixel size of 8.24 Å and a box size of 48 pixels, and subjected to three-dimensional classification into eight classes using a 30 Å low-pass-filtered initial model. The best-resolved class contained 423 particles and reached a resolution of 17.5 Å. This reconstruction was used as the template for the second iteration.

In the second iteration, template matching was performed on all 19 tomograms using the reconstruction obtained from the first iteration. The same cylindrical mask was used, and the template contrast was inverted to match that of the tomograms. This yielded 6,534 particles, which were extracted at bin 4 and classified into 10 three-dimensional classes. Selection of the best-resolved classes retained 4,279 particles. Further rounds of three-dimensional classification reduced the dataset first to 3,916 particles and then to 3,793 particles. Three-dimensional auto-refinement yielded a reconstruction at 17.8 Å resolution. The particles were subsequently re-extracted at bin 2 and refined to 17.7 Å resolution. Re-extraction at bin 1, followed by three-dimensional auto-refinement and post-processing using a 400-pixel mask, yielded a reconstruction at 9.7 Å resolution.

In the third iteration, the reconstruction obtained from the second iteration was used for template matching against the same 19 tomograms. A new mask was generated in RELION 5 from the input template, which was low-pass filtered to 16 Å and contrast-inverted before matching. This final template-matching round yielded 7,606 particles. Particles were extracted at bin 4 and classified into 10 three-dimensional classes. The selected set of 6,428 particles was re-extracted at bin 2 and subjected to further three-dimensional classification and two rounds of refinement to remove false-positive detections. The resulting set of 4,422 particles was re-extracted at bin 1 and refined, yielding a reconstruction at 14 Å resolution. Post-processing using a 400-pixel mask produced the final reconstruction at 9.4 Å resolution.

Local-resolution analysis was performed to assess spatial variation in map quality. The central branch-junction region, comprising Arp2, Arp3, ARPC1 and actin monomers adjacent to the junction, was resolved at higher resolution than the distal filament regions. The lower local resolution of the distal regions likely reflects filament flexibility and variation in filament length among subtomograms. Reduced local resolution around ARPC1 is consistent with previously reported conformational heterogeneity.

Previously reported atomic models of the Arp2/3-mediated branch junction (PDB 8P94) were rigid-body fitted into the final reconstruction using ChimeraX. The density resolved all seven Arp2/3 complex subunits together with adjacent actin monomers in the mother and daughter filaments. Additional densities compatible with cortactin and potentially phalloidin were observed but could not be assigned unambiguously at the available resolution.

Software used in this section was installed and configured by SBGrid (50).

#### Branch-junction spatial analysis

Three particle subsets were defined from the branch-junction dataset for downstream spatial analyses. The core subset comprised 1,004 branch junctions located within 11 manually defined podosome cores from seven tomograms and was used to analyze filament orientation relative to the ventral plasma membrane. The core-periphery subset comprised 726 branch junctions located within the outer 50% of the core radius in the same 11 podosomes and was used for radial-orientation analysis. The network-wide subset comprised 2,666 branch junctions from 17 tomograms, without separation of core and non-core regions, and was used for the Markov-chain analysis. Two tomograms in which cells extended into holes in the carbon support film were excluded from the network-wide analysis.

Particle metadata exported from RELION 5 were analyzed using a custom Python script (https://github.com/jasnin-lab/podosomes.git). For each tomogram, the analysis used the tomogram identifier, podosome-core origin, core radius and rotations required to align the ventral plasma membrane with the *x*–*y* plane. Particle coordinates and Euler angles were imported from the RELION particle STAR files, and the coordinates were transformed into a centered coordinate system in Å units.

For each tomogram, the ventral plasma membrane normal was calculated from the rotations required to align the membrane with the *x*–*y* plane. Reference vectors were transformed using standard right-handed rotation matrices, and the membrane normal was defined as their cross product. Particle orientations were calculated using the RELION ZYZ Euler-angle convention. In the branch-junction reference frame, the mother-and daughter-filament directions were defined as (1, 0, 0) and (cos 70*^◦^,* sin 70*^◦^,* 0), respectively, corresponding to the characteristic Arp2/3 branch angle. These vectors were transformed into the tomogram reference frame using the inverse particle rotation and used for subsequent orientation analyses.

For membrane-relative orientation analysis of the core and network-wide subsets, the signed angle between each mother-or daughter-filament direction vector and the ventral plasma membrane reference plane was calculated from normalized dot products and expressed in degrees. Angles were mapped to the range *−*90*^◦^* to +90*^◦^*, with 0*^◦^*corresponding to alignment with the membrane plane. Filaments with negative angles were classified as oriented toward the ventral plasma membrane and termed *downward*, whereas filaments with positive angles were classified as oriented away from the membrane and termed *upward*.

For radial-orientation analysis of the core-periphery subset, an inward-pointing displacement vector was defined from each particle coordinate to the corresponding podosome-core origin. Angles between this radial vector and the mother-or daughter-filament direction vectors were calculated both in three dimensions and after projection onto the *x*–*y* plane. In the projected analysis, daughter filaments forming angles below 90*^◦^* with the inward-pointing radial vector were classified as pointing toward the podosome center, whereas those forming angles above 90*^◦^* were classified as pointing away from the center.

### Hessian-based filament-orientation clustering

ActinSeg-derived F-actin segmentations from the whole-podosome and branch-junction datasets were used to determine point-wise filament orientations throughout the podosome volumes. For each actin voxel, we calculated the Hessian matrix of the segmented volume and defined the local filament orientation as the eigenvector associated with the minimum eigenvalue of the Hessian matrix. This yielded a non-polar director field in which the vectors **v** and *−***v** were treated as equivalent.

Point-wise orientation vectors from each podosome were clustered on the unit sphere using *K*-means clustering. Several values of *K* were evaluated, and *K* = 3 was selected because it reproducibly yielded three clusters of approximately equal size across tomograms, with a mean voxel proportion of 0.33 *±* 0.01 per cluster. Other values of *K* produced less consistent orientation classes across tomograms. The resulting clusters were defined as filament-orientation classes. For each orientation class *i*, voxel assignment was represented by the characteristic function *χ_i_*: R^3^ *→ {*0, 1*}*:

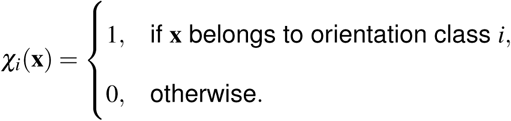

To generate local orientation domains from the point-wise measurements, each characteristic function *χ_i_*was convolved with a normalized uniform three-dimensional kernel of dimensions 3 nm *×* 21 nm *×* 21 nm. The analysis was restricted to the convex hull of the segmented F-actin network, which defined the podosome volume used for subsequent orientation-domain and layered-architecture analyses. For each orientation class *i*, the convolved field was denoted by *g_i_*, and the local class probability was calculated as:

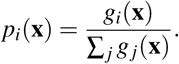

The dominant orientation class at each voxel was defined as the class with the highest local probability.

Angular separations between orientation classes were calculated as non-polar angular distances between their corresponding mean orientation vectors. The mean inter-class angular separations were 61*^◦^ ±* 8*^◦^*and 66*^◦^ ±* 13*^◦^* in the whole-podosome and branch-junction datasets, respectively. Within-class angular dispersion was quantified as the angular distance between each point-wise filament-orientation vector and the mean direction of its assigned class. The mean within-class angular dispersions were 36*^◦^ ±* 23*^◦^* and 37*^◦^ ±* 23*^◦^* in the whole-podosome and branch-junction datasets, respectively. Together, these measurements defined three principal axes of filament orientation.

To determine whether the filament-orientation classes formed same-orientation layers along the membrane-normal axis, dominant class assignments were analyzed as a function of axial position relative to the ventral plasma membrane. For each podosome, the *z*-axis was defined as normal to the local membrane plane. Same-orientation layer thickness was measured along this axis as the distance over which a given dominant orientation class persisted before transitioning to another class. The mean layer thicknesses were 11.3 *±* 5.0 nm and 11.7 *±* 5.1 nm in the whole-podosome and branch-junction datasets, respectively.

### Markov-chain analysis of Arp2/3-mediated branching

For the branch-junction dataset, branch-junction positions and orientations obtained by template matching and subtomogram averaging were mapped onto the ActinSeg-derived filament segmentations. For each Arp2/3-mediated branch junction, local mother-and daughter-filament orientations were assigned from the refined branch-junction orientation and the surrounding filament-orientation field.

A three-component Gaussian mixture model partitioned daughter-filament orientations relative to the ventral plasma membrane into three dominant states, which we termed *downward*, *parallel* and *upward* daughter-filament orientation states. These fixed boundaries were then applied to the daughter-filament angle distributions in each 15-nm height band. Daughter-state distributions were similar across bands along the membrane-normal axis, consistent with a common transition framework throughout the network. We therefore modeled branching as a Markov chain in which the probability distribution of the daughter-filament state depends only on the orientation state of the corresponding mother filament. Paired mother-and daughter-state assignments were used to construct a 3 *×* 3 transition matrix. Each matrix entry represents the conditional probability that a mother filament in a given orientation state produces a daughter filament in a specified orientation state. The matrix was row-normalized so that the transition probabilities from each mother-filament state summed to one.

Eigendecomposition of the network-wide transition matrix identified a unique stationary distribution comprising 27% *downward*, 54% *parallel* and 19% *upward* daughter-filament states (Fig. 5c). Because membrane-proximal layers have accumulated later during network growth, the lower half of each podosome was treated as the structurally more advanced region. We therefore quantified daughter-filament orientation states in this region and compared their distribution with the stationary distribution derived from the network-wide transition matrix. The resulting proportions were 28% downward, 51% parallel and 21% upward, closely matching the stationary distribution.

To assess variation in daughter-filament orientation-state distributions along the membrane-normal axis, height was defined as the distance from the ventral plasma membrane along the local membrane-normal direction. Each podosome volume was divided into successive 15-nm height bands, and branch junctions were assigned to bands according to their height above the membrane. Daughter-filament angular distributions were calculated separately for each height band. Applying the previously fitted Gaussian mixture model to each height-resolved distribution yielded the empirical fractions of downward, parallel and upward daughter filaments.

The resulting orientation-state vector **f***_k_* for the *k*-th height band was expressed as a linear combination of the normalized eigenvectors of the network-wide transition matrix:

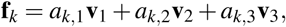

where **v**_1_, **v**_2_ and **v**_3_ are the normalized eigenvectors of the 3 *×* 3 transition matrix, with **v**_1_ corresponding to the stationary eigenvector, and *a_k,_*_1_, *a_k,_*_2_ and *a_k,_*_3_ are the corresponding real-valued expansion coefficients for the *k*-th height band.

For visualization, the absolute values of the expansion coefficients were used to represent the relative contributions of the three eigenvectors. For each height band, these coefficient magnitudes were normalized by dividing each magnitude by their sum:

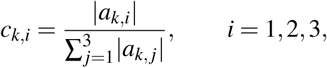

such that

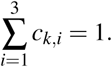

For clarity in Fig. 5d, the height-dependent normalized coefficient magnitudes *c_k,_*_1_, *c_k,_*_2_ and *c_k,_*_3_ are labeled *c*_1_, *c*_2_ and *c*_3_, respectively.

### Layered-helical analysis and assembly pseudo-age inference

Axial recurrence of filament-orientation classes was quantified from the spatial autocorrelation of each orientation-class probability along the membrane-normal axis. To minimize boundary effects and improve numerical stability, each tomogram was divided into a 3 *×* 3 grid in the plane parallel to the ventral plasma membrane, and the analysis was restricted to the central region. For orientation class *i*, the autocorrelation as a function of axial displacement *z* was calculated as:

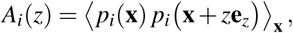

where **e***_z_* is the unit vector normal to the ventral plasma membrane and the average was calculated over positions **x** within the analyzed volume. In a periodically organized field, recurrent structural features produce local autocorrelation maxima at displacements corresponding to their axial spacing. This approach enabled recurrence spacing to be measured without explicitly segmenting the undulating same-orientation layers.

The resulting autocorrelation profiles decayed non-monotonically with increasing axial displacement. Local maxima beyond the zero-offset peak marked recurrent occurrences of the same orientation class, and the distances between successive maxima were used to determine same-orientation layer spacing. This analysis yielded mean spacings of 30.9 *±* 6.6 nm and 34.9 *±* 5.2 nm in the whole-podosome and branch-junction datasets, respectively.

To assess the helical repeat, filament orientations were treated as non-polar directors, such that orientations differing by 180*^◦^* were considered equivalent. In the whole-podosome dataset, the accumulated angular rotation over one three-class cycle was estimated as 3 *×* 61.4*^◦^* = 184.2*^◦^*. Because this value differs by only approximately 4*^◦^* from the equivalent 180*^◦^*non-polar orientation, the orientation pattern returned to an approximately equivalent state after one complete cycle. The effective axial repeat of the layered-helical organization was therefore defined by the recurrence spacing of the same orientation class along the membrane-normal axis.

The progressive accumulation of recurrent orientation layers was used as a structural indicator of podosome assembly state. For each podosome, the pseudo-age was defined as 1 plus the rounded mean number of nonzero-offset local maxima detected across the autocorrelation profiles of the three orientation classes:

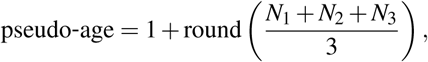

where *N_i_* is the number of nonzero-offset local maxima detected for orientation class *i*. The additional value of 1 accounts for the initial layer represented by the zero-offset autocorrelation peak. This procedure provides a discrete structural metric based on the recurrence of orientation classes rather than on absolute podosome height, thereby reducing sensitivity to local membrane curvature and force-induced axial deformations.

## Data availability

Representative tomograms, ActinSeg-derived filament segmentations, and subtomogram averages of F-actin and Arp2/3-mediated branch junctions generated in this study will be deposited in the Electron Microscopy Public Image Archive (EMPIAR) and the Electron Microscopy Data Bank (EMDB) before publication. Accession codes will be provided upon deposition. Processed data supporting the analyses, including filament-coordinate lists, polarity assignments, Arp2/3 branch-junction coordinates and orientations, filament-orientation-class assignments, Markov transition matrices and figure source data, will be provided as supplementary data. All other data supporting the findings of this study are available from the corresponding authors upon reasonable request.

## Code availability

ActinSeg is available at https://github.com/jasnin-lab/actinseg.git. Custom scripts used for filament resampling, polarity assignment, podosome geometric analysis, Arp2/3 branch-junction spatial analysis, Hessian-based filament-orientation analysis, layered-helical analysis, Markov-chain modeling and assembly pseudo-age inference will be made available at https://github.com/jasnin-lab/podosomes.git before publication. Third-party software and version information are provided in the Methods.

## Acknowledgements

We acknowledge Vanessa Soldan from the METi imaging facility (Genotoul-TRI), a member of the national infrastructure France-BioImaging supported by the French National Research Agency (ANR-24-INBS-0005, FBI BIOGEN). We thank Christophe Thibault from LAAS-CNRS for technical assistance. We are grateful to Martin Lenz for helpful discussions on the modeling, and to Thomas Schwarz-Romond and members of the Cryoskeleton Lab for constructive comments on the manuscript. This work was supported by the Agence Nationale de la Recherche (ANR) and the Deutsche Forschungsgemeinschaft (DFG) through the ANR–DFG NLE 2020 grant JA-3038/2-1 awarded to R.P. and M.J., and by financial support from ITMO Cancer of Aviesan, with funds administered by Inserm, awarded to R.P. J.V.O. received support from the Helmholtz Association through the joint research school “Munich School for Data Science – MUDS”.

## Author contributions statement

R.P. and M.J. conceived, designed and supervised the project. J.R.-B. prepared primary human macrophages and performed immunofluorescence, fluorescence imaging, podosome size analysis and protrusion force microscopy. S.B. performed cell unroofing and vitrification on EM grids. J.S. and M.J. acquired the cryo-ET data, and J.S. and T.D. reconstructed the tomograms. J.S. performed F-actin segmentation in Amira, and D.O. performed ActinSeg-based F-actin segmentation. J.S. designed and performed the F-actin subtomogram analysis, polarity assignment and podosome geometric analysis. T.D. performed the branch-junction spatial analysis. J.V.O. designed and performed the Hessian-based filament-orientation clustering and layered-helical analysis. J.V.O. designed the Markov-chain analysis of Arp2/3-mediated branching, which was performed by T.D. and J.V.O. J.V.O. performed the assembly pseudo-age analysis. M.J. supervised and interpreted all cryo-ET analyses. J.S., J.R.-B., T.D., J.V.O., D.O., R.P. and M.J. prepared the figures. R.P. and M.J. acquired funding. M.J. wrote the manuscript with input from all authors. All authors reviewed and approved the manuscript.

