## Supplementary Information for "Layered helical order reveals assembly states in podosome actin networks"

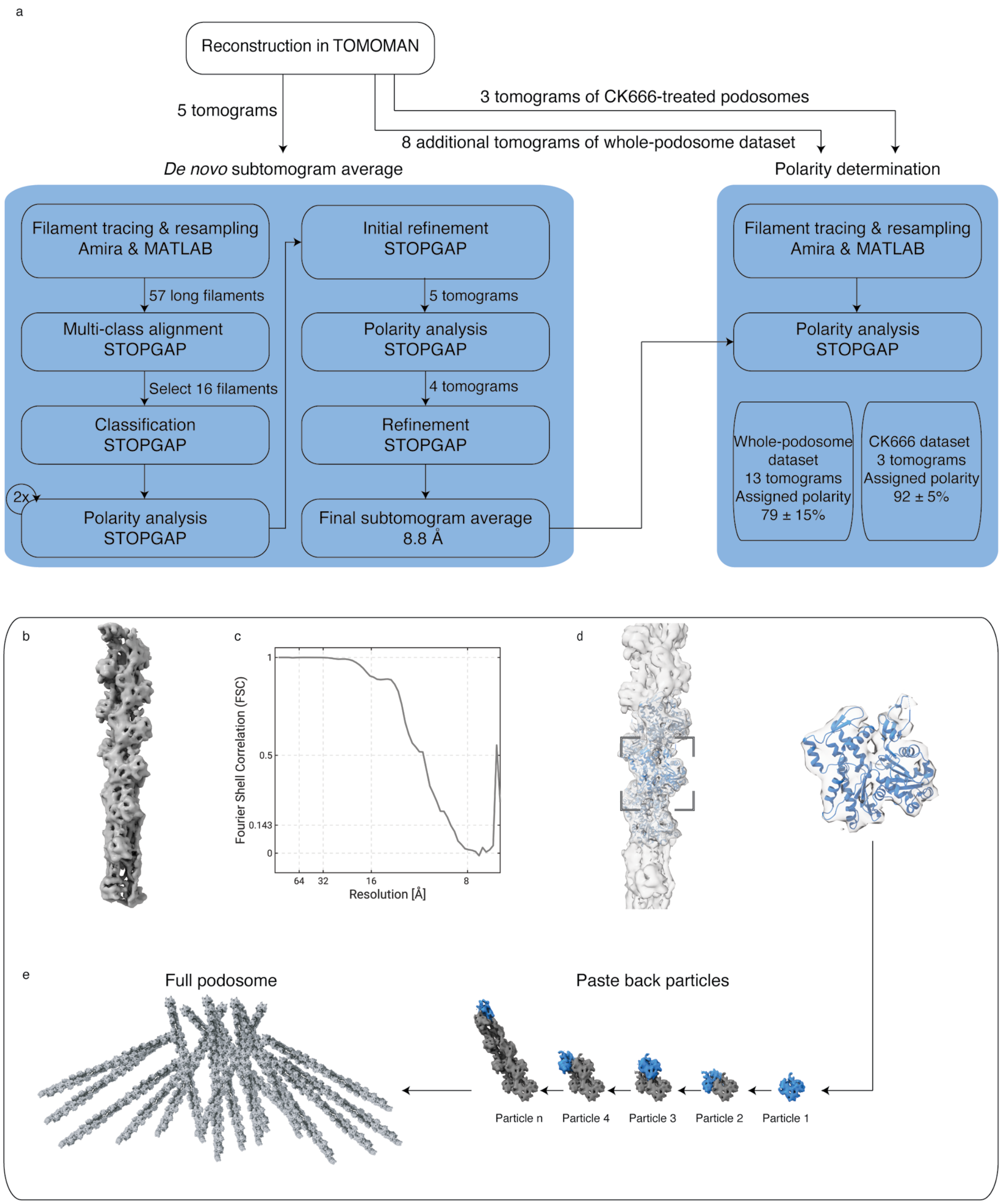


**Figure S1. Subnanometer F-actin structure and polarity-assignment workflow in human macrophage podosomes. a,** Workflow for *de novo* F-actin subtomogram averaging and filament-polarity assignment. An initial reference generated from five whole-podosome tomograms was refined in STOPGAP and used to assign polarity in the whole-podosome and CK666-treated datasets. Polarity was assigned to 93 $\pm$ 5% of filaments in the CK666-treated dataset. **b,** Final F-actin subtomogram average at 8.8 Å resolution. **c,** Fourier shell correlation curve; the dashed line indicates the 0.143 criterion. **d,** Atomic F-actin model fitted into the subtomogram average, with the boxed region enlarged on the right. **e,** Schematic of polarity propagation from refined actin-monomer positions to complete filaments and the podosome network. Blue highlights mark the same actin-monomer region across successive particles.

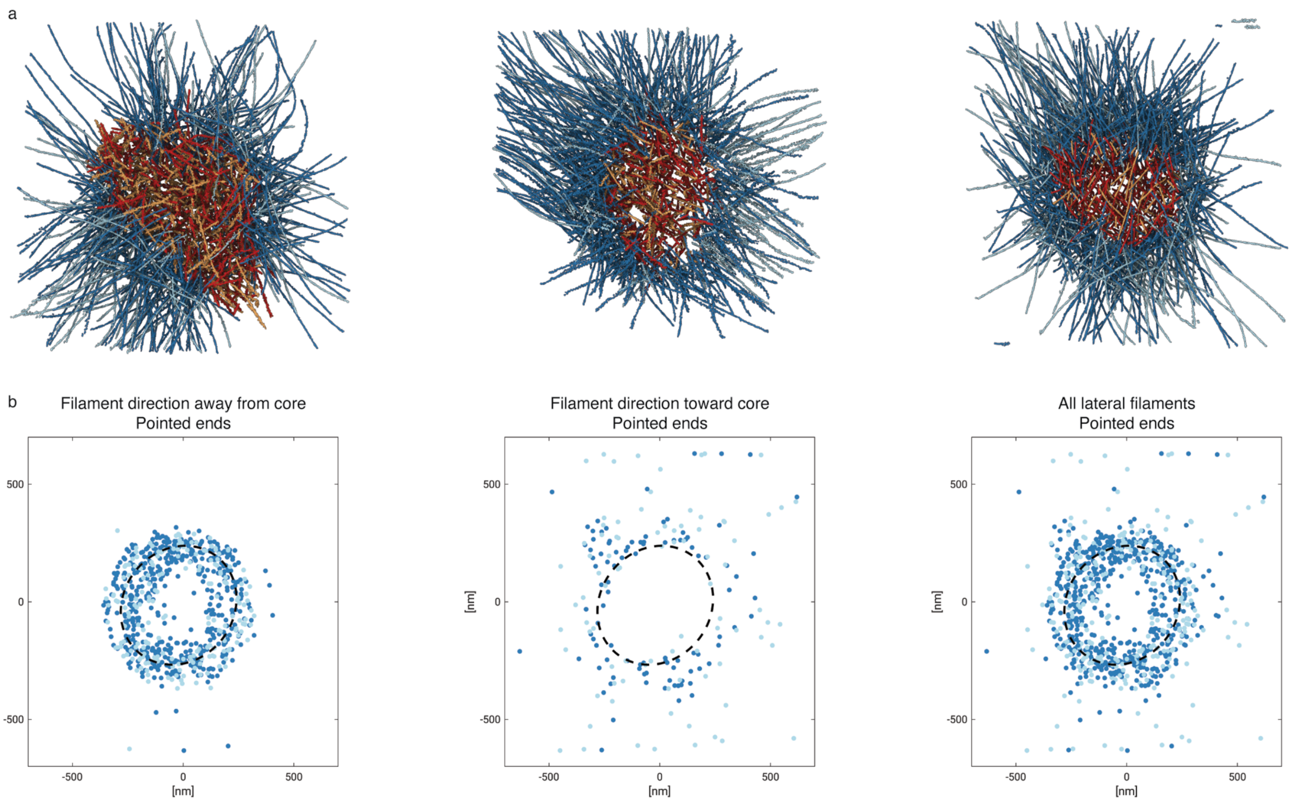


**Figure S2. F-actin polarity analysis across human macrophage podosomes. a,** Top views of polarity-resolved F-actin networks in podosomes #1, #2 and #7. Core filaments are colored according to whether their barbed ends are directed toward (red) or away from (orange) the ventral plasma membrane (VM); lateral filaments are shown in dark and light blue, respectively. **b,** Spatial distributions of lateral-filament pointed ends in podosome #3. From left to right, plots show outward-directed, inward-directed and all lateral filaments. Colors follow the convention in **a**. Dashed circles mark the core boundary.

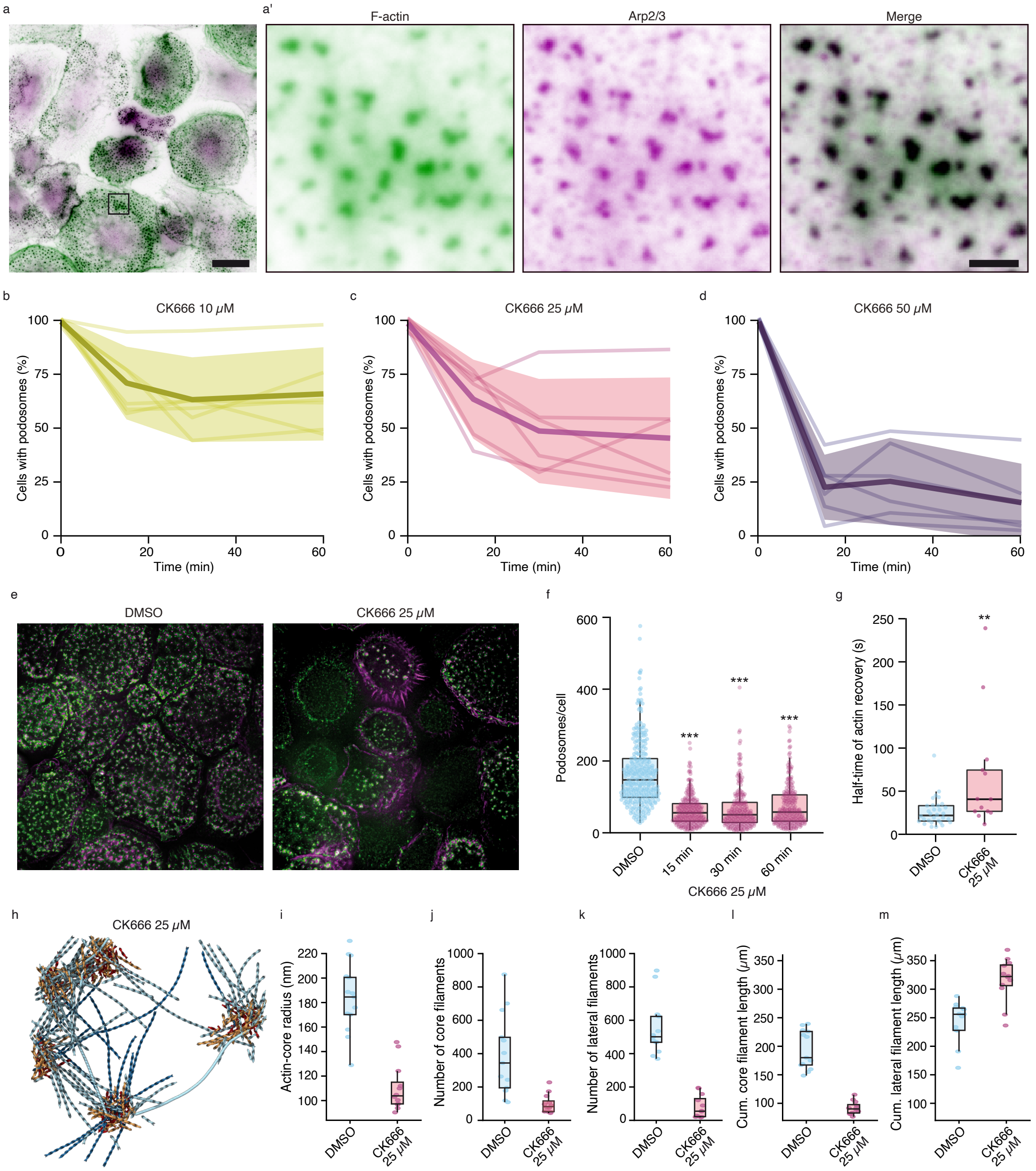


**Figure S3. Partial Arp2/3 inhibition impairs podosome assembly and actin-network organization. a,a',** Localization of F-actin and Arp2/3 in human macrophage podosomes. The boxed region in **a** is enlarged in **a'**. Scale bars, 20 $\mu$m and 2 $\mu$m, respectively. **b–d,** Fraction of macrophages containing podosomes during treatment with 10 $\mu$M (**b**), 25 $\mu$M (**c**) or 50 $\mu$M (**d**) CK666. Lines represent three donors; shaded regions show mean $\pm$ SD. **e,** Representative control and 25 $\mu$M CK666-treated macrophages. F-actin is magenta and vinculin green. Scale bar, 20 $\mu$m. **f,** Podosome number per cell during treatment with 25 $\mu$M CK666. $n = 3$ donors and $n > 200$ cells per condition. Kruskal–Wallis test with Dunn's multiple-comparison test; ***$P < 0.001$. **g,** Actin-recovery half-time at podosome cores under control and 25 $\mu$M CK666-treated conditions. $n = 3$ donors and $n > 24$ cells per condition. Two-tailed Mann–Whitney test; **$P < 0.01$. **h,** Polarity-resolved F-actin network in CK666-treated podosome #1. **i–m,** Cryo-ET quantification of actin-core radius (**i**), core- and lateral-filament numbers (**j,k**) and cumulative filament lengths (**l,m**). Boxes show the median and interquartile range, with 10th–90th percentile whiskers. The analysis includes 13 control podosomes from 13 tomograms and 12 CK666-treated podosomes from 3 tomograms.

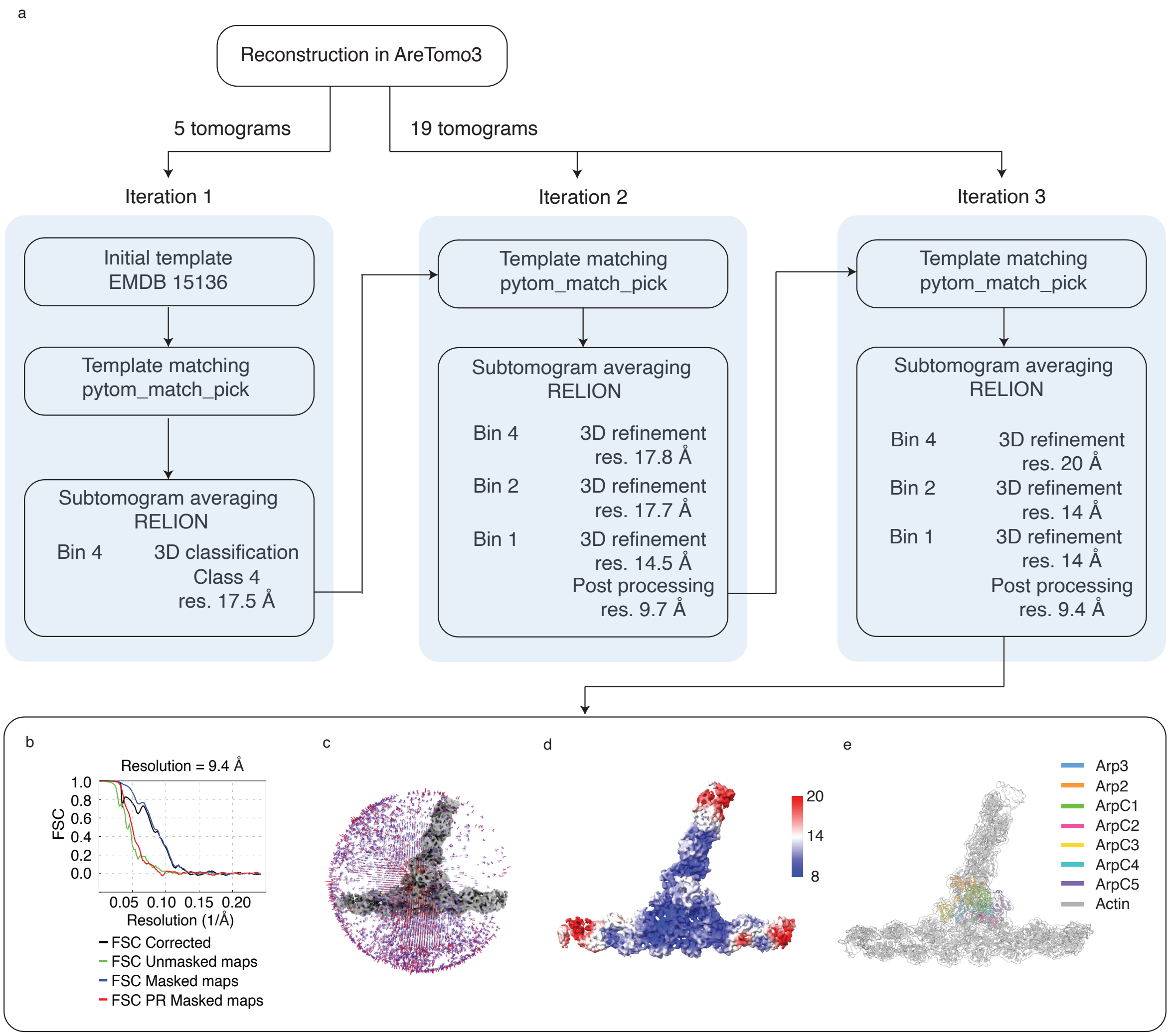


**Figure S4. Detection and subnanometer reconstruction of Arp2/3-mediated branch junctions. a,** Workflow for branch-junction detection and reconstruction. Three iterative rounds of template matching in PyTom and subtomogram averaging in RELION 5 yielded a final reconstruction at 9.4 Å resolution from 19 tomograms. **b,** Fourier shell correlation curves; resolution was determined using the 0.143 criterion. **c,** Angular distribution of particles included in the final reconstruction. **d,** Local-resolution map. **e,** Rigid-body fitting of the seven Arp2/3 subunits and adjacent mother- and daughter-filament actin monomers.

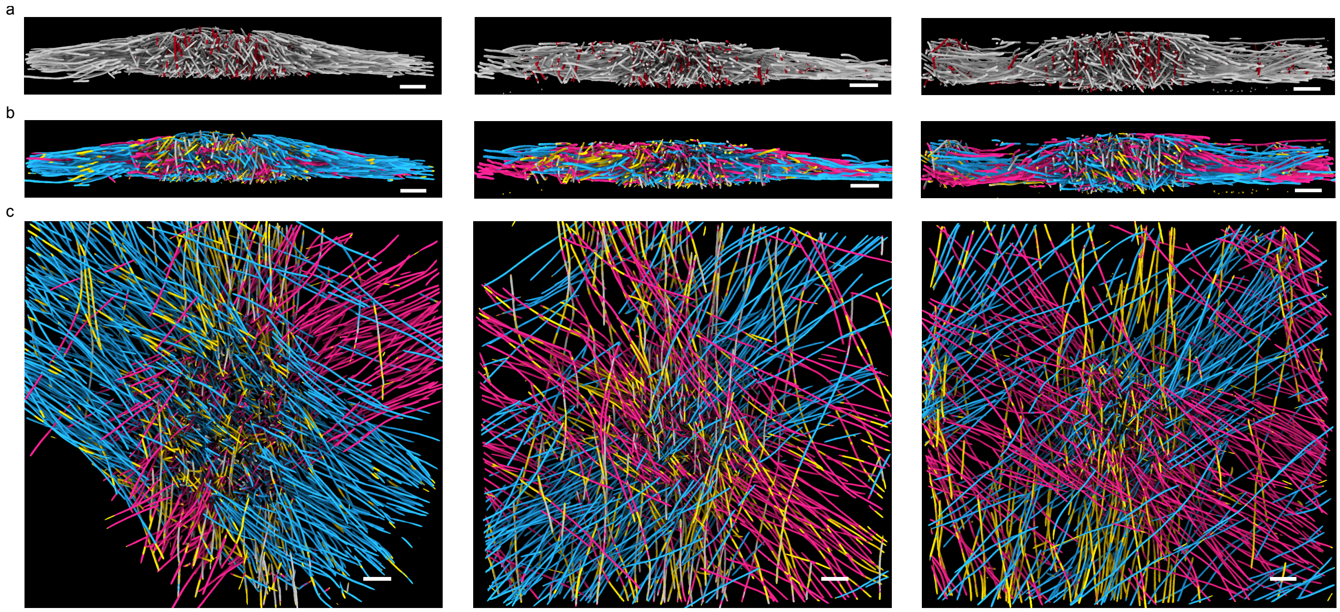


**Figure S5. Structural architecture and filament orientation of podosomes. a,** Side views of ActinSeg-derived F-actin segmentations from three representative podosomes. Excluded upright core filaments are shown in red. **b,** Corresponding side-view orientation maps illustrating the layered organization of the three filament-orientation classes. **c,** Corresponding top views illustrating the spatial distribution and principal directions of the orientation classes. Podosomes correspond to #2, #4 and #5, from left to right, respectively. Scale bars, 100 nm.

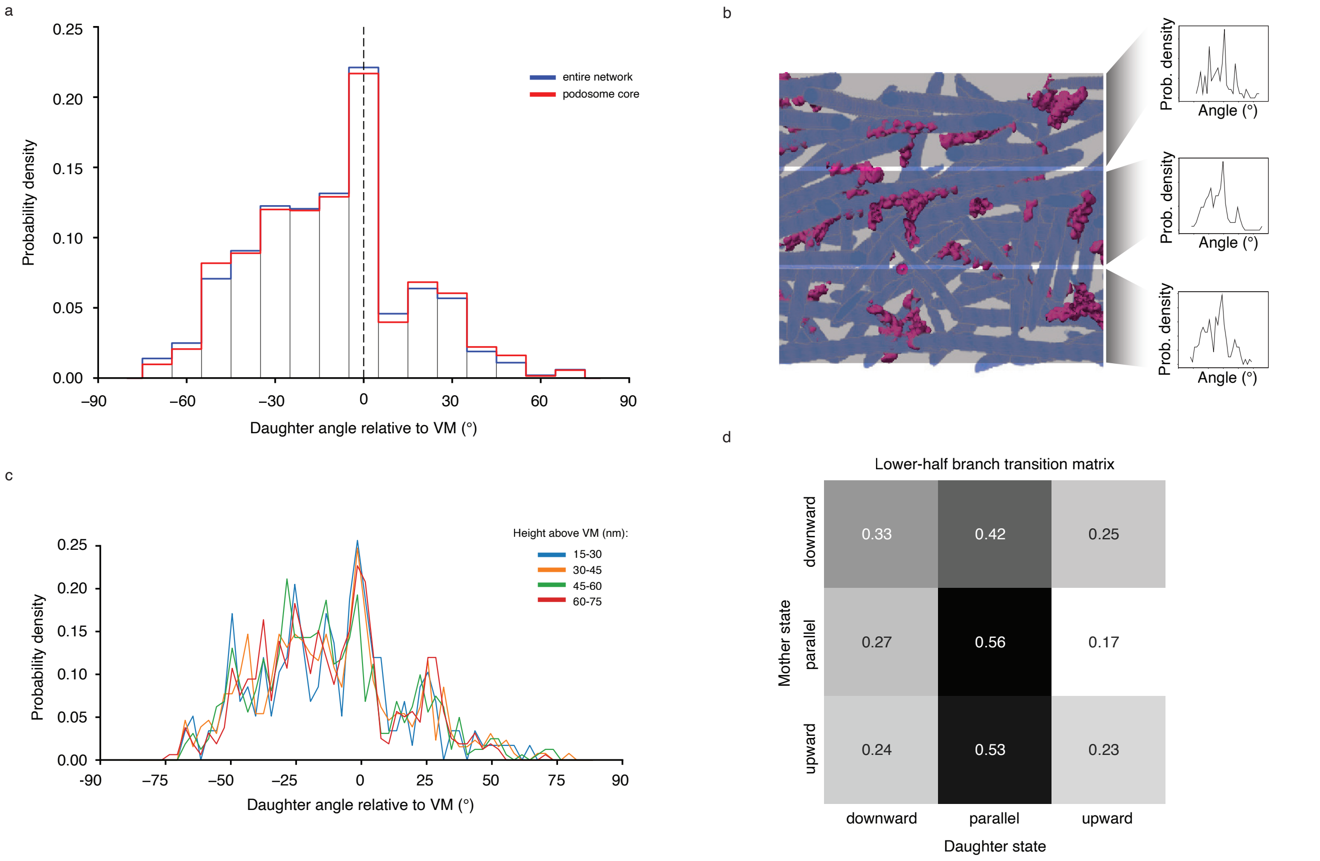


**Figure S6. Arp2/3-mediated transitions between branch-orientation states. a,** Daughter-filament orientation distributions for the network-wide subset and the core subset, comprising 17 tomograms and 11 podosome cores from 7 tomograms, respectively. Angles are measured relative to the ventral plasma membrane (VM). **b,** Assignment of branch junctions to successive 15-nm height bands along the membrane-normal axis. **c,** Daughter-filament orientation distributions within successive height bands in the network-wide subset. **d,** Mother-to-daughter orientation-state transition matrix for branch junctions in the lower half of podosomes.

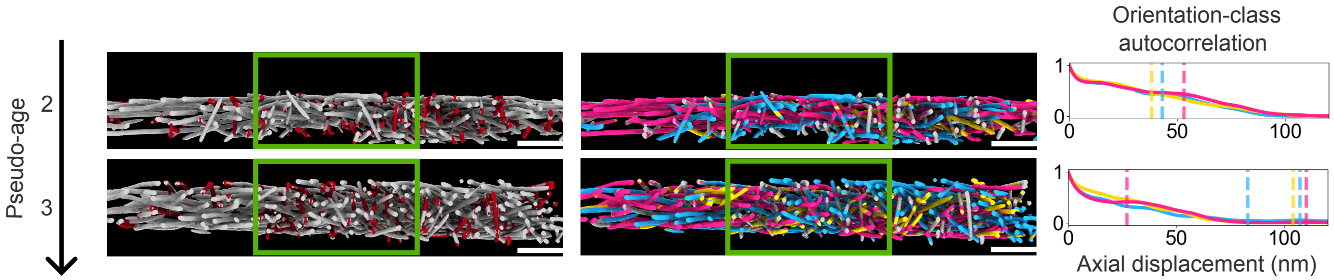


**Figure S7. Pseudo-age estimation of podosomes based on layered structural order.** Representative F-actin segmentations (left), filament-orientation maps (middle) and orientation-class autocorrelation profiles (right) for podosomes with increasing pseudo-age. Green boxes indicate the analyzed central regions. Colored curves represent the three orientation classes, and vertical lines mark the nonzero-offset autocorrelation maxima used to calculate pseudo-age. Podosomes from the branch-junction dataset correspond to #25 and #21, from top to bottom. Scale bars, 100 nm.
